# Chaperone-mediated heterotypic phase separation regulates liquid-to-solid phase transitions into amyloid fibrils

**DOI:** 10.1101/2024.06.13.598862

**Authors:** Sandeep K. Rai, Roopali Khanna, Anusha Sarbahi, Ashish Joshi, Samrat Mukhopadhyay

## Abstract

Biomolecular condensates formed via the phase separation of proteins and nucleic acids are thought to regulate a myriad of cellular processes with exquisite spatiotemporal precision. However, such highly dynamic, viscoelastic, mesoscopic, intracellular membraneless bodies can undergo aberrant liquid-to-solid transitions into a range of amyloid-like species. The formation of such pathological assemblies necessitates their clearance by the cellular protein quality control machinery comprising molecular chaperones. Nonetheless, the mechanism underlying the chaperone-mediated regulation of protein homeostasis within biomolecular condensates remains elusive. Here, we present a unique case demonstrating that a heat shock protein 40 (Hsp40), Ydj1, promotes the heterotypic phase separation of intrinsically disordered tau via intermolecular electrostatic and hydrophobic interactions. Through a diverse array of tools involving high-resolution fluorescence imaging, single-droplet steady-state and picosecond time-resolved fluorescence anisotropy, and single-molecule FRET (Förster resonance energy transfer), we elucidate the diverse structural conformations of tau present within phase-separated heterotypic condensates that are otherwise predisposed to aggregation. Our vibrational Raman spectroscopy and electron microscopy data show that the presence of Ydj1 in tau-Ydj1 condensates abolishes the formation of amyloid fibrils, unlike tau-only droplets. By sequentially deleting segments, we identify amyloidogenic hexapeptide motifs located in the hydrophobic microtubule-binding region of tau that foster contacts with the peptide-binding regions of Ydj1, promoting the formation of tau-Ydj1 binary condensates. Additionally, we show that the underlying network of interactions governing these condensates can be further tuned by RNA. Our results underscore an intriguing interplay of molecular drivers that govern chaperone-associated phase separation, with broader implications for the chaperoning of a wide range of intrinsically disordered proteins involved in physiology and disease.

## Introduction

Biomolecular condensate formation via macromolecular phase separation of proteins and nucleic acids offers effective spatiotemporal organization and subcellular localization of cellular components into highly dynamic, noncanonical, regulable membraneless organelles (1–15). These phase-separated membraneless compartments are now known to orchestrate a plethora of cellular events ranging from biochemical reactions, cellular transport, and signaling to genome processing, nuclear organization, and so forth. The formation of these intracellular membraneless bodies is governed by a multitude of weak, transient, multivalent, and homotypic and heterotypic interactions between proteins and nucleic acids. Intrinsically disordered proteins/regions (IDPs/IDRs) and proteins comprising oligomerization domains are the key candidates for macromolecular phase separation due to their ability to introduce multivalency into a system via the formation of ephemeral noncovalent contacts such as electrostatic, hydrophobic, cation-π, and π-π interactions (4,9,12,14–18). These intermolecular interactions result in the formation of viscoelastic complex fluids via a range of mechanisms involving percolation-coupled density transitions, coupled associative and segregative phase transitions, and so forth (4,5). The physical properties of such phase-separated condensates are highly tunable under various external and internal stimuli such as small molecules, nucleic acids, and post-translational modifications (PTMs) (19–26). Any perturbations in these tightly regulated membraneless bodies in intracellular space can result in changes in their material properties, consequently disrupting their physiological functions, as implicated in a range of neurodegenerative diseases, including amyotrophic lateral sclerosis, frontotemporal lobar dementia, etc. (26–31).

Microtubule-associated tau is a neuronal IDP that plays an essential role in maintaining the cytoskeletal network of cells and mediating transport in neurons (32–35). As a result of its inherent sequence attributes, tau undergoes biomolecular condensation (36–41). Functional tau-rich condensates can undergo stress-mediated aberrant phase transitions associated with a change in their material state to form pathological amyloid-like aggregates, which are the hallmarks of neurodegeneration involved in an array of tauopathies (42). A growing body of research has revealed the broad role of molecular chaperones that form the cellular protein quality control (PQC) network in modulating the phase behavior and several pathological tau species (43–49). Molecular chaperones, including heat shock proteins (Hsps), encompass a group of highly-conserved, related families of proteins that are crucial for maintaining cellular stability via protein homeostasis, with Hsp40 being an essential member of such multi-chaperone complex-controlled pathways (50–53). The ATP-independent Hsp40 family of proteins, typified by a characteristic Hsp70-binding J-domain, acts as the first line of defense against misfolded proteinaceous species in cells. These J-domain proteins are associated with a highly-conserved domain architecture across species that includes an N-terminal J-domain, a glycine-phenylalanine (G/F) linker, a zinc-finger-like region, and peptide binding C-terminal domains (CTD I/II), followed by a dimerization-domain (50, 51). Despite the emphasis placed on the importance of the Hsp40-mediated chaperoning of tau and its significance, both in terms of tau pathology and function, the underlying molecular mechanism by which the cellular network modulates condensate properties remains elusive. Here, we present a unique case of the modulation of the phase behavior of tau by Ydj1, a yeast homolog of the human class-I Hsp40, DnaJA1. Using a diverse array of methodologies that allow us to access a wide range of spatiotemporal resolutions, we unmasked the role of the tau sequence grammar in governing its interaction with Ydj1 and illuminated the crucial molecular events associated with the conformational shapeshifting of tau.

## Results

### Ydj1 promotes phase separation of tau

Tau is a polyampholytic IDP comprising clustered oppositely charged residues in its distinct domains, enabling it to undergo phase separation mediated by interdomain electrostatic interactions (Fig. 1*A*, S1*A*). As a prelude, we started by performing phase separation assays of tau under our conditions at a physiological pH, as demonstrated previously by us and others (Fig. 1*B-D*) (36–41, 54). To probe the phase behavior of the yeast Hsp40, Ydj1, used for our experiments (Fig. 1*E*), we first established the structural homology between Ydj1 and human Hsp40s by comparing the structure of Ydj1 with that of its conventionally used human analog. On comparing their structures, we observed a considerable similarity in their domain architectures (Fig. 1*F*). Their similarity is also supported by previous studies that have reported the mechanistic basis of their action and robustness through different species (55–57). To test the behavior of Ydj1, we performed confocal imaging of Ydj1 (10 µM) in the presence of ∼ 1% of Ydj1 labeled with Alexa Fluor 488-C5-maleimide and visually confirmed its inability to phase separate on its own (Fig. 1*G*). After establishing the absence of Ydj1 condensation, we set out to elucidate its effect on the phase behavior of tau by performing turbidity measurements in the presence of Ydj1 (Fig. 1*H*). The turbidity of tau (10 µM) sharply increased indicating that Ydj1 promotes tau phase separation. Next, we estimated the saturation concentration (C_sat_) of tau in the absence and the presence of Ydj1 and found it to be lower in the latter case further indicating Ydj1-mediated potentiation of tau phase separation (Fig. 1*I*). To directly observe and verify the co-phase separation of tau and Ydj1, we introduced a single cysteine at the 244^th^ position (tau Q244C) on a null-cysteine variant of full-length tau (C291S, C322S) via site-directed mutagenesis and labeled it using Alexa Fluor 594-C5-maleimide. To visualize tau-Ydj1 droplets, we doped unlabeled tau (10 µM) and Ydj1 (10 µM) in the reaction mixture (in 20 mM HEPES, pH 7.4) with ∼ 1% of each of their corresponding labeled counterparts and performed two-color confocal Airyscan imaging (Fig. 1*J*, S1*B)*. Our imaging results showed a colocalization of tau and Ydj1 in droplets, which we further corroborated by the Pearsons’ coefficient colocalization and Manders’ overlap analysis (Fig. 1*K*). These droplets underwent rapid fusion, acquired a spherical shape, and showed a rapid fluorescence recovery after photobleaching (FRAP), indicating their dynamic liquid-like nature (Fig. 1*L*). In addition, our FRAP measurements for Ydj1 in tau-Ydj1 condensates showed a somewhat slower recovery compared to tau suggesting the formation of a denser network formed by the heterotypic association of tau and Ydj1. Our fluorescence correlation spectroscopy (FCS) experiments further supported this observation and exhibited a slower diffusion time for Ydj1 compared to tau in the heterotypic mixture (Fig. 1*M, N*, and S1*C*). Overall, this set of results demonstrated the Ydj1-mediated potentiation of tau phase separation resulting in the heterotypic tau-Ydj1 condensates comprising a denser network formed by the two proteins. Next, to discern the underlying molecular drivers governing tau-Ydj1 co-phase separation, we aimed to probe the effect of ionic strength and temperature on these heterotypic condensates.

**Fig. 1.**
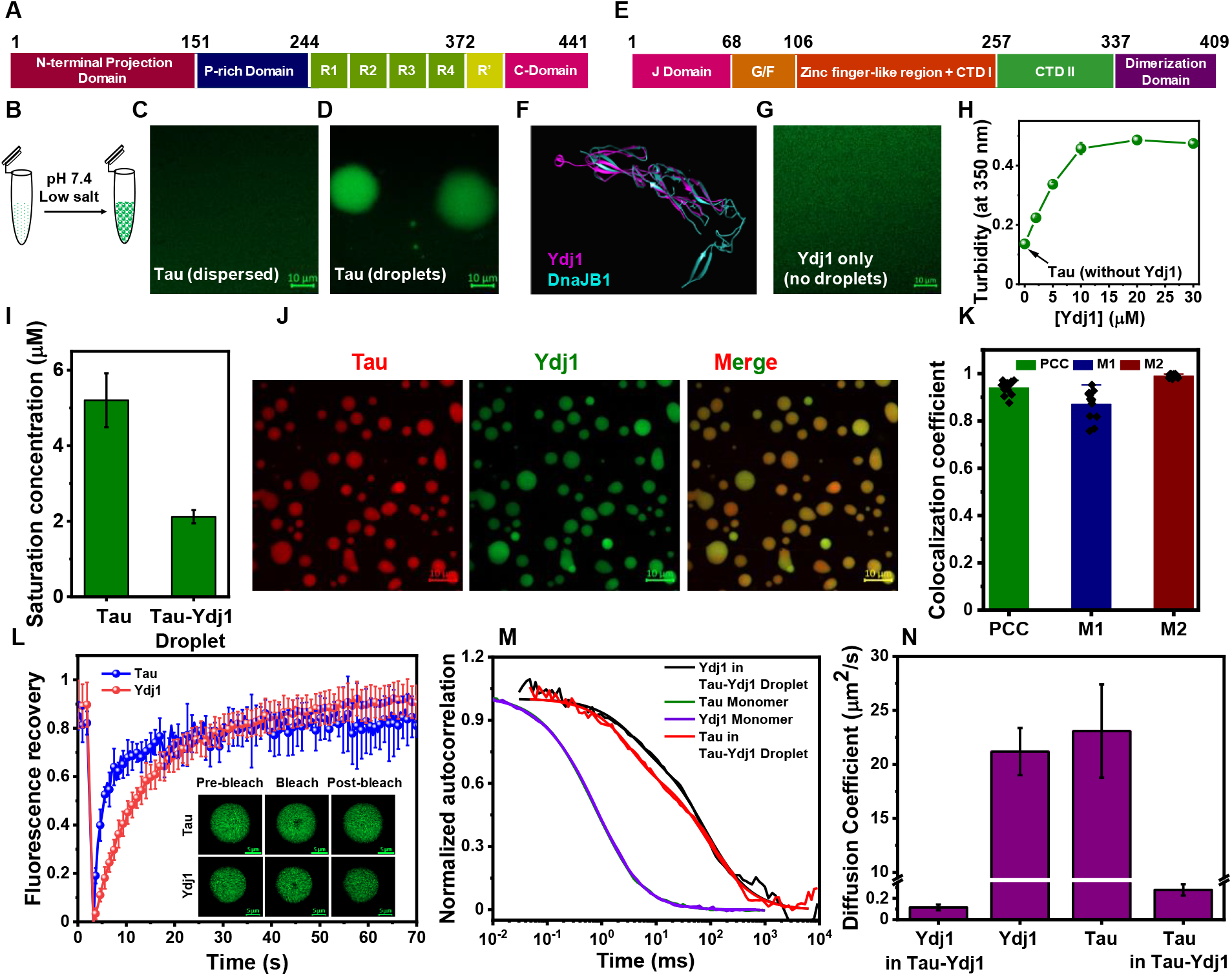
Tau phase separation is modulated by Ydj1. (*A*) Domain architecture of the full-length human tau protein. We used a null cysteine variant (C291S, C322S) of the same for our studies. For the amino acid sequence of tau, see Fig. S1*A.* (*B*) Tau phase separation can be induced at low salt and physiological pH conditions. (*C*) Tau (tau-Q244C-Alexa Fluor 488, green) remains in the monomeric dispersed phase in the presence of salt. (*D*) Tau condensates were reconstituted using purified protein (10 µM) in the absence of salt at physiological pH (20 mM HEPES, 2 mM DTT, pH 7.4; Scale bar, 10 μm). (*E*) Domain architecture of Ydj1, a yeast analog of human DnaJA1, which is a member of the Hsp40 family of molecular chaperones. (*F*) The yeast and human variants are homologous, as seen by the structural alignment of Ydj1 with another J-domain containing Hsp40 (Cyan: DnajB1; PDB: 2QLD; Magenta: Ydj1; PDB: 1NLT; RMSD between atom pairs = 1.030 Å). Here, due to the absence of a solved structure of DnaJA1, the human homolog of Ydj1, the next available Hsp40 member, DnaJB1, was used for structure comparison. (*G*) Ydj1 (10 µM unlabeled, doped with sparsely labeled Ydj1-Alexa Fluor 488, green) does not phase separate alone under our conditions (Scale bar, 10 μm). (*H*) Turbidity (O.D. at 350 nm) measurements of tau-Ydj1 reaction mixtures as a function of increasing Ydj1 concentration. The concentration of tau was kept constant at 10 μM. The data represent mean ± SD; n = 3. (*I*) The saturation concentration (C_sat_) of tau for tau-only and tau-Ydj1 condensates was estimated by high-speed centrifugation. (*J*) A high-resolution two-color Airyscan confocal image of colocalized tau (tau-Q244C-Alexa Fluor 594, red) and Ydj1 (sparsely labeled with Alexa Fluor 488, green) in tau-Ydj1 condensates (Scale bar, 10 μm). Both tau and Ydj1 concentrations were 10 μM. (*K*) Plots of Pearson’s and MandeR′s colocalization coefficients quantifying the presence of tau in the droplets of Ydj1 and vice-versa (The data represent mean ± SD; n = 15 high-resolution confocal images). (*L*) FRAP kinetics of tau-Ydj1 droplets. Alexa Fluor 488-labeled proteins were used for both tau and Ydj1 for independent FRAP studies. The data represent mean ± SD; n = 5. The inset shows fluorescent images acquired during FRAP measurements (Scale bar, 5 μm). (*M*) The normalized autocorrelation plot was obtained from FCS measurements performed for monomeric dispersed tau and Ydj1, as well as for tau and Ydj1 in tau-Ydj1 droplets. The unnormalized autocorrelation plots are shown in *Supporting Information*, Fig. S1. (*N*) Diffusion coefficients of Ydj1 in tau-Ydj1 condensates, Ydj1 in monomeric dispersed form, and tau in monomeric dispersed form and tau-Ydj1 droplets obtained from FCS measurements for five different droplets. For all FCS measurements of monomers, 5 nM of labeled proteins were used (tau-Q244C-Alexa Fluor 488, Ydj1 sparsely labeled with the same dye). For droplets, 0.5 nM of labeled proteins were used in both cases. Unlabeled tau and Ydj1 concentrations were both 10 μM. See Supporting Information for details.

### An interplay of electrostatic and hydrophobic interactions regulates the complex phase separation of tau and Ydj1

Heterotypic phase separation resulting in a network of biomolecules is generally facilitated by a combination of enthalpic as well as entropic driving forces (39, 58–60). Considering the prevalence of hydrophobic residues as well as charged residues in the sequence space of both tau and Ydj1, we first examined the contributions of hydrophobic and electrostatic interactions in their co-phase separation. As previously established, the ability of Hsps, including Ydj1, to interact with non-native or misfolded forms of proteins may depend on the formation of hydrophobic contacts via its peptide binding region (PBR) (61). Therefore, we started by setting out to discern the role of increasing temperature on tau-Ydj1 condensation. Our turbidity and microscopy experiments suggested a temperature-dependent increase of tau-Ydj1 condensation indicative of a lower critical solution transition (LCST) behavior (Fig. 2*A*, S2*A*). In general, purely LCST behavior is typical for phase separation in systems driven by hydrophobic effects involving the entropic release of water molecules. A change in the co-phase separation propensity of the two proteins, as observed by our turbidity measurements in the presence of increasing concentrations of 1,6-hexanediol, further highlights the importance of hydrophobic interactions in the tau-Ydj1 system (Fig. S2*B*). At a physiological pH, Ydj1 exists as a negatively charged polypeptide (pI = 6.30), while tau exhibits a net positive charge under the same condition (pI = 8.24) (Fig. S2*C*, S2*D*) (62). The oppositely charged nature of these two proteins suggests an important role of electrostatic interactions in this system. To test this, we performed two-color confocal imaging at increasing salt concentrations. In agreement with our hypothesis, droplet formation was abrogated at higher salt concentrations underscoring the importance of electrostatic interactions in the co-phase separation of tau and Ydj1 (Fig. 2*B*). Next, we asked if alteration in the net charge on tau due to phosphorylation could influence the tau phase behavior.

**Fig. 2.**
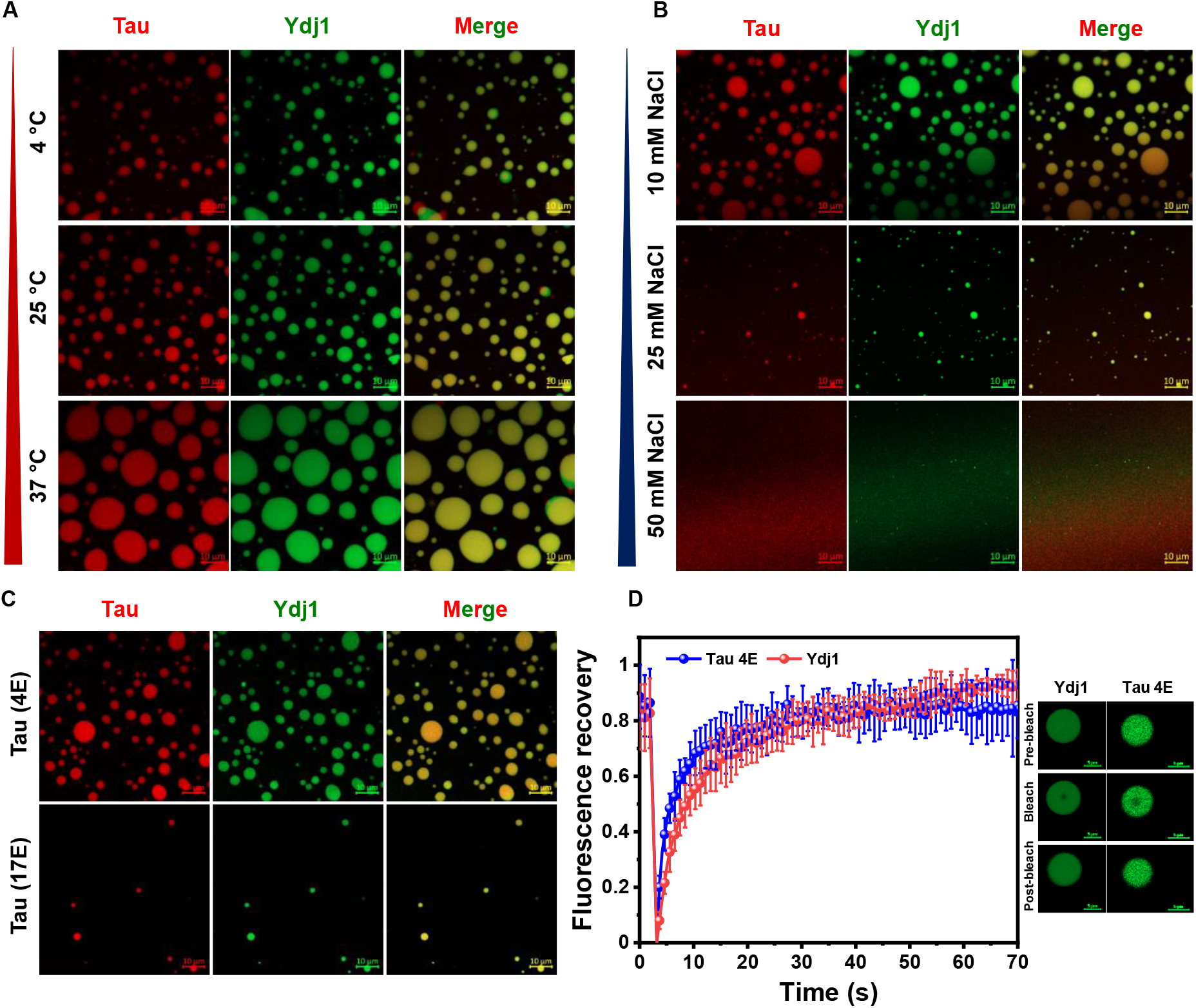
An interplay of electrostatic and hydrophobic interactions drives tau-Ydj1 condensation. Two-color Airyscan confocal images of tau (10 μM, red) and Ydj1 (10 μM, green) heterotypic condensates (*A*) at increasing temperatures and (*B*) in the presence of an increasing concentration of NaCl (Scale bar, 10 μm). (*C*) Tau 4E (10 μM, red) and Ydj1 (10 μM, green) droplets (*Upper*) compared with droplets formed by tau 17E and Ydj1 (10 μM, green, *Lower*; Scale bar, 10 μm). (*D*) FRAP kinetics of tau 4E-Ydj1 droplets. Alexa Fluor 488-labeled proteins were used for both tau and Ydj1 for independent FRAP studies (Both tau and Ydj1 concentrations were 10 μM). The data represent mean ± SD; n = 5.

PTMs of tau such as phosphorylation by specific kinases are pathologically significant. They are also associated with changes in the charge profile of tau giving rise to altered intra- and interdomain interactions in tau. Moreover, previous studies have also highlighted phosphorylation-associated changes in the phase behavior of tau (33, 36, 63). To recapitulate the same in our system, we used phosphomimetic tau mutants with phosphorylation patterns similar to those achieved by cellular kinases. One of these mutants (tau 4E; pI =7.16), where four residues were mutated to glutamate, recapitulated the phosphorylation usually attained by microtubule affinity-regulating kinases (36). The second mutant (tau 17E; pI = 5.60) was created by selectively replacing serine, threonine, and tyrosine residues throughout the tau chain to glutamate, as achieved by protein kinase A that phosphorylates tau to a significantly greater extent (64). Upon performing our turbidity and imaging experiments with these mutants in the presence of Ydj1, we observed that although the tau 4E phase separated with Ydj1 (Fig. 2*C*, *D*, and S2*E*), the greater negative charge on tau 17E abolished this interaction presumably due to the repulsion between the two protein chains under our condition (Fig. 2*C*, S2*E*). Altering the electrostatic interactions using phosphomimetic mutants further highlighted the importance of electrostatic interactions in modulating tau-Ydj1 co-phase separation. Taken together, our salt- and temperature-dependent data, coupled with experiments using phosphomimetic tau mutants, suggested an intricate interplay of both electrostatic and hydrophobic interactions in driving tau-Ydj1 co-phase separation. Next, to further dissect the domain-specific interactions of this heterotypic co-condensation, we performed our experiments with truncated variants of tau.

### Central and hydrophobic regions of tau are responsible for tau-Ydj1 co-condensation

After establishing the change in the phase behavior of tau in the presence of Ydj1, we set out to discern the role of different domains of tau that are crucial for heterotypic interactions. To this end, we performed our experiments with biologically relevant truncations of tau (Fig. 3*A*). We began by using a negatively charged N-terminal fragment of tau (Nh2htau; residues 26-230; pI = 5.32) comprising the projection and the proline-rich domains. This truncation is involved in the pathology of several tauopathies and is also found in amyloid-like deposits in the brains of diseased patients (65). As expected, we observed no droplet formation by this mutant in the presence of the similarly-charged Ydj1 (Fig. 3*B*) due to the repulsion between the two protein chains. Inspired by studies that have hinted at the importance of the tau-repeat region in its interaction with various members of distinct chaperone families, we next used the well-studied tau K18 fragment, which comprises only the four oligopeptide repeats from R1 to R4 (Fig. 3*A*) (33, 46). In the presence of Ydj1, we observed droplet formation by tau K18 as suggested by our turbidity measurements and imaging, albeit at much higher protein concentrations (Fig. 3*C*, S3*A*, S3*B*). Moreover, the droplets formed were much smaller than those formed in the presence of full-length tau, even at higher protein concentrations. We further performed our experiments with a positively-charged C-terminal fragment of tau (tau truncation, 151-391, pI = 10.23) that comprises the proline-rich region repeats P1 and P2 in addition to the central (R1-R4) and pseudo repeat regions (R′) (Fig. 3*A*) (66). In contrast to Nh2htau, our confocal imaging and turbidity measurements with tau truncation showed the formation of droplets, much like full-length tau, even at much lower protein concentrations (Fig. 3*D*, S3*C*). Our data suggested the importance of the repeat region as well as the neighboring positively charged proline-rich region of tau in driving tau-Ydj1 coacervation. This further validates our salt and temperature-dependent measurements in unraveling the importance of an interplay of electrostatic and hydrophobic interactions in the formation of tau-Ydj1 condensates. In addition to being essential for microtubule binding, the central repeat containing MTBR of tau also contains two-hexapeptide motifs, PHF6* (^275^VQIINK^280^) and PHF6 (^306^VQIVYK^311^) at the beginning of the second and the third repeats, respectively, which are crucial for tau fibrillation (Fig. 3*A*) (33). To assess the importance of these aggregation-promoting hexapeptide motifs in tau-Ydj1 phase separation, we created two deletion mutants, namely tauΔPHF6* and tauΔPHF6, where we selectively deleted one hexapeptide while keeping the other. Under our phase-separating condition, while tauΔPHF6* showed a phase behavior similar to full-length tau, the propensity of tauΔPHF6 to undergo phase separation was considerably lower in the presence of Ydj1 as suggested by a lower solution turbidity and sparse droplet formation in our imaging experiments (Fig. 3*E*-*H*, S3*D*). This observation further highlighted the importance of the hydrophobic hexapeptide stretch, PHF6, in tau-Ydj1 co-condensation. Additionally, by using the N and C-terminal truncated variants of Ydj1 (Ydj1 N-terminal; residues 1-207, and Ydj1 C-terminal incorporating hydrophobic CTDs I and II; residues 208-409), we showed that the C-terminal region of Ydj1, which is also known as the peptide binding region, is essential for its interaction and co-phase separation with tau (Fig. S3*E-H*).

**Fig. 3.**
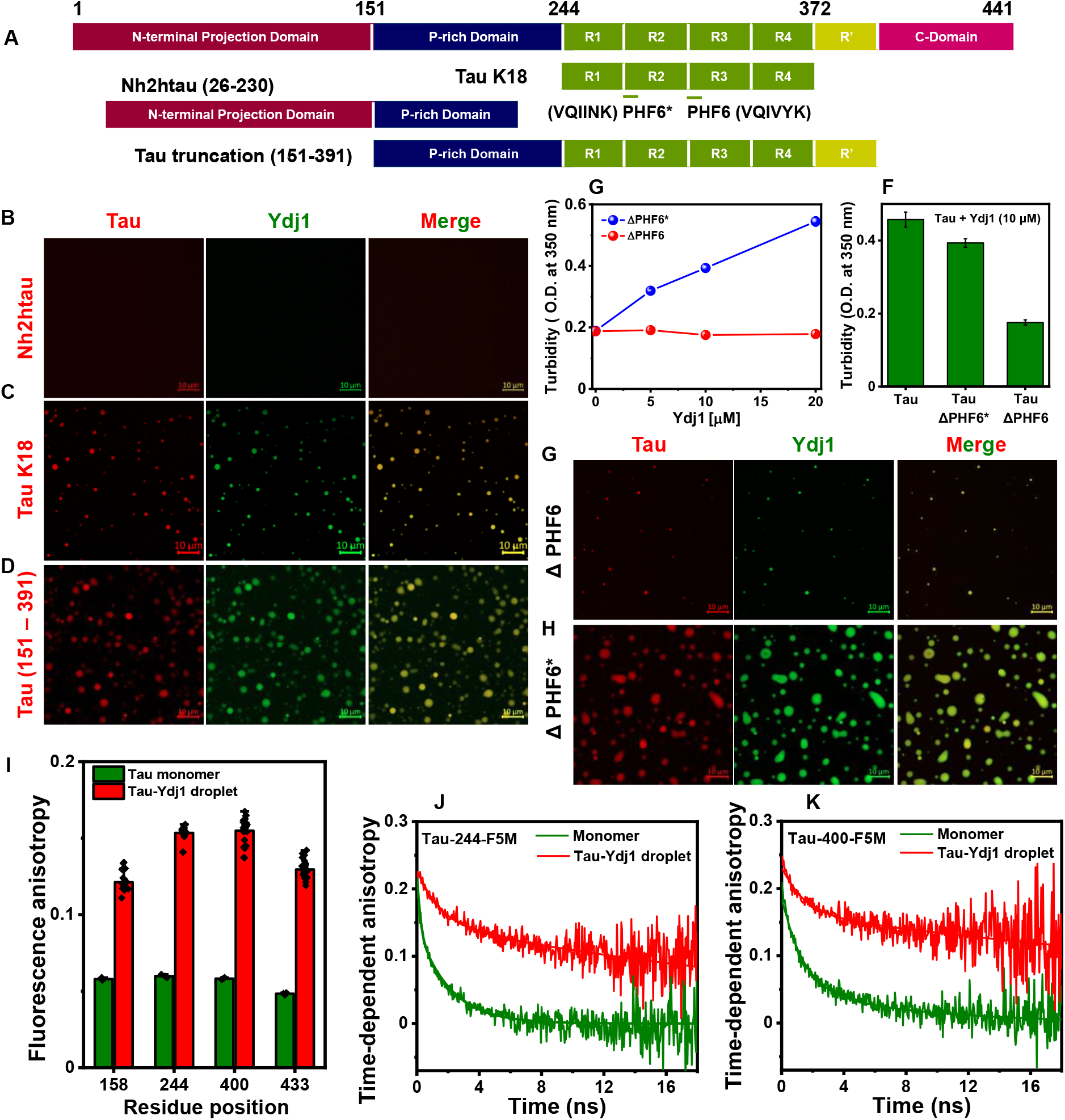
Domain-specific interactions are fundamental to tau-Ydj1 phase separation. (*A*) Depiction of the tau constructs used. Two-color Airyscan confocal images of (*B*) Nh2htau (10 μM, red) and Ydj1 (10 μM, green) (Scale bar, 10 μm), (C) tau K18 (10 μM, red) and Ydj1 (10 μM, green) (Scale bar, 10 μm) and, (*D*) tau truncation (10 μM, red) and Ydj1 (10 μM, green) (Scale bar, 10 μm). (*E*) Turbidity plots of tauΔPHF6 (10 μM) and tauΔPHF6* (10 μM) as a function of the Ydj1 concentration. The data represent mean ± SD; n = 3. (*F*) Comparison of turbidity (O.D. at 350 nm) of droplet reactions of full-length tau (10 μM), tauΔPHF6 (10 μM), and tauΔPHF6* (10 μM) in the presence of Ydj1 (10 μM). The data represent mean ± SD; n = 3. Two-color Airyscan confocal images of (*G*) tauΔPHF6 (10 μM, red) and Ydj1 (10 μM, green) droplets compared to (*H*) tauΔPHF6* (Scale bar, 10 μm). (*I*) Single-droplet steady-state fluorescence anisotropy measurements of F5M labeled single-Cys mutants of tau spanning the sequence in the dispersed, monomeric state (olive) and in tau-Ydj1 droplets (red). Data for more than n = 30 different droplets were considered for droplet anisotropy. For data for other F5M labeled single-Cys mutants of tau, see *Supporting Information*, Fig. S3*D*. (*J*) Time-resolved fluorescence anisotropy decays of F5M labeled tau Q244C and (*K*) tau S400C of tau in monomeric state (olive) and tau-Ydj1 droplets (red). The solid lines are fits obtained using decay analysis. Tau and Ydj1 concentrations were both 10 μM. See *Supporting Information, Methods* for details of measurements and analysis and *Supporting Information*, Table S2 for recovered parameters.

Next, to investigate the regio-specific conformational changes of tau upon tau-Ydj1 condensation, we performed site-specific single droplet steady-state fluorescence anisotropy measurements using fluorescein-5-maleimide (F5M)-labeled single cysteine variants of tau encompassing the entire tau polypeptide chain. Such site-specific fluorescence anisotropy measurements allowed us to determine the local conformational mobility. Our single droplet steady-state anisotropy measurements exhibited a sharp increase in the anisotropy values upon condensation suggesting an increase in the ordering and the presence of a dense rotationally restricted environment within tau-Ydj1 condensates. We observed the highest anisotropy values for the residues lying in the MTBR indicating its greater structural ordering and the formation of preferable contacts with Ydj1 facilitated by this region (Fig. 3*I*, S3*I*, and S3*J*). Since steady-state anisotropy provides only time-averaged information, we performed single-droplet picosecond time-resolved fluorescence anisotropy measurements to temporally discern the different modes of rotational dynamics exhibited by the tau protein chain within tau-Ydj1 condensates. Here, we chose residues lying in the repeat region of tau, i.e., the 244th and 400th positions in the tau sequence. Our single-droplet measurements showed that tau exhibited slower depolarization kinetics (global rotational correlation time ∼ 40 ns) compared to monomeric tau (Fig. 3*J*, *K*, and Table S2). Such an increase in the slower rotational correlation time indicated the formation of a dense network within tau-Ydj1 co-condensates impeding the reorientation dynamics of the polypeptide chain. Having established the domains and regio-specific factors governing tau-Ydj1 droplet formation, we next investigated the RNA-mediated modulation of these co-condensates. Because of its ability to act as a significant cofactor modulating tau liquid-to-solid transitions, RNA can have a strong impact on tau-Ydj1 phase behavior.

### Competing interactions in the tau-Ydj1-RNA ternary system govern the properties of the multi-component condensates

In cells, phase-separated condensates exist as complex assemblies of biomacromolecules such as proteins and nucleic acids. The polymeric nature of the constituents of these condensates affords multivalency to the system allowing the formation of a multi-component network (21–24). In biological systems, the physical properties of nucleic acids such as RNA also contribute to the nature of compartmentalization achieved in addition to serving other more traditionally well-studied roles. The charge on any given RNA chain, in addition to its ability to recruit multiple protein partners and introduce multivalency into a system, makes RNA a potent scaffold in numerous biological condensates (67). To test the effect of RNA on our system, we performed our experiments in the presence of increasing concentrations of polyU RNA (Fig. 4*A*, *B*, and S4*A*). Within a concentration range from 5-40 ng/µL, we observed no change in the morphology of the droplets compared to those formed in the absence of RNA (Fig. 4*B*). Our FRAP experiments also showed high recovery in the presence of RNA indicative of the liquid-like nature of these condensates. Interestingly, although both tau and Ydj1 achieved complete recovery, the recovery of Ydj1 was faster than what we observed for droplets formed with just the addition of the protein components (Fig. 4*C*, *D*). The faster diffusion of Ydj1 in the presence of RNA suggested an overall relaxation of Ydj1 in the condensed phase via RNA-mediated multivalency and competing interactions between the negatively charged Ydj1 and RNA for tau in the tau-Ydj1-RNA ternary system. At higher concentrations of RNA, we observed the complete dissolution of condensates as expected for typical RNA-associated reentrant phase behavior (Fig. 4*A*, *B*) (68–70). Additionally, to visualize and monitor the RNA dynamics within coacervates, we labeled the 5’ end of RNA with the Alexa Fluor 488-NHS ester (succinimidyl ester) dye via the EDC-NHS-coupling mediated activation of its 5′ phosphate (Fig. S4*B*) (71). Our three-color Airyscan confocal imaging experiments confirmed the localization of RNA with the protein components in tau-Ydj1 condensates (Fig. 4*E*). Next, in order to estimate the internal mobility of RNA within these condensates, we performed FRAP experiments using labeled RNA. These results revealed a slower fluorescence recovery in comparison to tau and Ydj1 suggesting the formation of a dense RNA-mediated network in the interior of the heterotypic condensates (Fig. 4*C*). In addition, our results are in line with previous reports on the dynamics of RNA in droplets (72). Since the central region of tau establishes contacts with Ydj1 and acts as the primary site for RNA interaction owing to the presence of lysine clusters (73), we set out to delineate the role of RNA in modulating tau truncation-Ydj1 coacervates. Our multi-color imaging experiments captured a change in the morphology of the condensates formed by this ternary system, in contrast to the previously observed completely mixed condensates observed for tau truncation-Ydj1 (Fig. S4*C*). In the presence of RNA, these condensates adopted a vacuolar morphology, as reported previously for non-stoichiometric multi-component mixtures, which undergo dissolution at high concentrations of RNA (74). Additionally, in the three-color imaging performed for this system, we observed the colocalization of all three components of the mixture in the rims of the resulting vacuolar condensates (Fig. S4*D*). This set of results indicated that an interplay of competing interactions between tau, Ydj1, and RNA controls the physical properties of ternary condensates. Since protein-rich condensates are known to act as reaction crucibles for aberrant liquid-to-solid phase transitions, we next asked whether Ydj1 could play an important role in arresting such condensate maturation into amyloid-like aggregates.

**Fig. 4.**
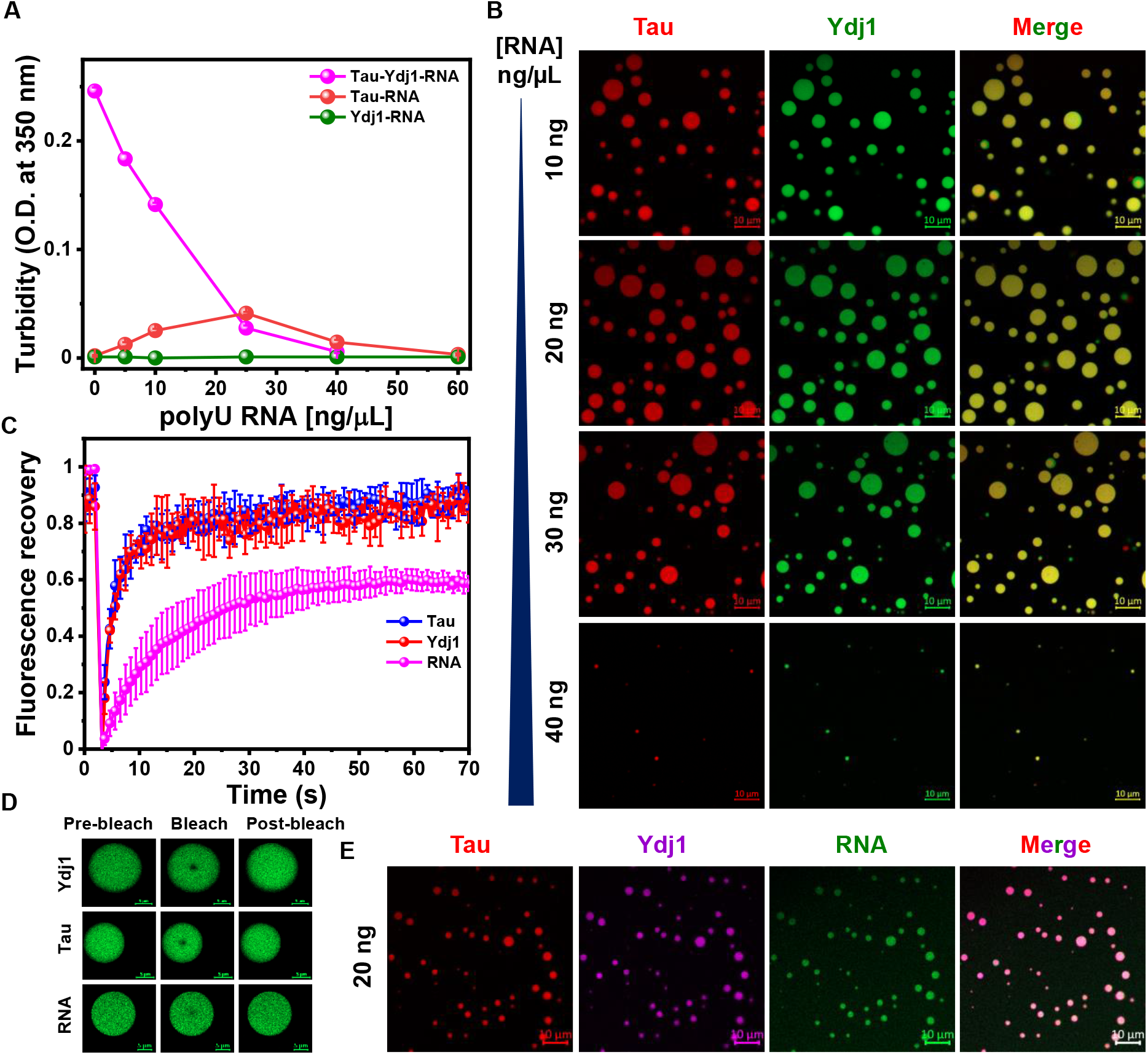
RNA-mediated Tau-Ydj1 reentrant phase behavior. (*A*) Turbidity plots of tau-Ydj1-RNA, tau-RNA, and Ydj1-RNA as a function of RNA concentration (both tau and Ydj1 concentrations were 10 μM). The data represent mean ± SD; n = 3. (*B*) Two-color Airyscan confocal images of the tau-Ydj1-RNA (tau-red; Ydj1-green) ternary system as a function of increasing RNA concentrations (Scale bar, 10 μm). (*C*) Three-color Airyscan confocal image of colocalized tau (tau-244C-Alexa Fluor 594, red); Ydj1 (Alexa Fluor 647, pink) and RNA (5’ Phosphate-Alexa Fluor 488, green) in tau-Ydj1-RNA coacervates (Scale bar, 10 μm). Both tau and Ydj1 concentrations were 10 μM, while RNA concentration was 20 ng/μL. (*D*) FRAP kinetics of tau-Ydj1-RNA droplets. The data represent mean ± SD; n = 5. (*E*) Fluorescence images of droplets during FRAP measurements (Scale bar, 5 μm). Alexa Fluor 488-labeled proteins and RNA were used for Ydj1 (*Upper*), tau (*Middle*), and RNA (*Lower*) for independent FRAP studies (Both tau and Ydj1 concentrations were 10 μM, and RNA concentration was 20 ng/μL).

### Ydj1 halts liquid-to-solid phase transitions and aggregation of tau

The highly concentrated environment within biomolecular condensates makes them particularly susceptible to aberrant phase transitions. This is especially true for several neuronal IDPs because of their ability to engage in multivalent interactions within such condensates, as previously observed for polypeptides such as FUS, TDP-43, and tau (26–31). Prior evidence has confirmed the conversion of monomeric tau into amyloid-like aggregates, both in the presence as well as in the absence of cofactors, which is accelerated several folds upon phase separation in the presence of cofactors or crowding. Several such potentially pathogenic events in cells are promptly halted or counteracted by the cellular chaperone machinery comprising several Hsps (43–48, 75). To recapitulate this phenomenon in our system in a crowder or cofactor-free environment, we incubated the tau-Ydj1 reaction mixtures with agitation and monitored the aggregation kinetics using thioflavin T (ThT) which is a well-known amyloid marker. As expected, over time, phase-separated tau exhibited ThT fluorescence intensity over ∼ 4 µM protein concentration suggesting the formation of amyloid-like aggregates in a nucleation-dependent manner (Fig. 5*A*, S5*A*). This was further corroborated by the electron microscopy imaging of the end products, where we observed the formation of fibrillar species (Fig. 5*B*). Monomeric tau, on the other hand, did not aggregate under the monitored conditions. In the presence of Ydj1, however, we saw sub-stoichiometric ratio-dependent inhibition of tau aggregation, and in the presence of 10 µM. No measurable increase in the ThT fluorescence intensity was observed indicating the Ydj1-mediated abrogation of tau aggregation. The final species of tau-Ydj1 reaction mixtures contained non-amyloid protein conglomerates (Fig. 5*A*, *B*) even at sub-stoichiometric concentrations of Ydj1 (Fig. S5*B*). Our observation is in line with previous reports where tau was shown to form such higher-order non-amyloid species in the presence of a complex of two major molecular chaperones, namely, Hsp70 and Hsp90 (46,76). Hence, instead of halting tau in its monomeric state, tau-Ydj1 condensation leads to the formation of nanoscopic hetero-clusters of tau in complex with Ydj1. To further characterize the nature of these species, we used vibrational Raman spectroscopy and monitored the amide I vibrational band (1620-1720 cm^-1^), which originates primarily due to the C = O stretching of the polypeptide backbone, assigned to the secondary structure of the protein (Fig. 5*C*). Our Raman experiments showed that the shape and the width of the amide I Raman band remained unchanged indicating the structural arresting of the tau-Ydj1 condensates (Fig. 5*C*, 5*D*). In contrast, upon aging, tau condensates in the absence of Ydj1, exhibited a much sharper amide I band having maxima around ∼ 1675 cm^-1^ which is a hallmark of amyloid cross-β architecture (Fig. 5*E*, 5*F*). Additionally, our ThT kinetics with the tau-Ydj1-RNA ternary system also showed similar aggregation kinetics as the binary system in the absence of RNA, signifying the importance of Ydj1 in modulating the liquid-to-solid phase transition of tau even in the presence of cofactors like RNA, which are typically associated with aggregation (Fig. S5*C*). Taken together, this set of results showed that Ydj1 could serve as a potent regulator of tau solidification and amyloid formation. Next, to elucidate the mechanistic basis of the Ydj1-mediated inhibition of tau fibrillation and to discern the key events associated with conformational changes, we set out to perform single-molecule FRET experiments that are capable of recording conformational distributions and subpopulations in a molecule-by-molecule manner.

**Fig. 5.**
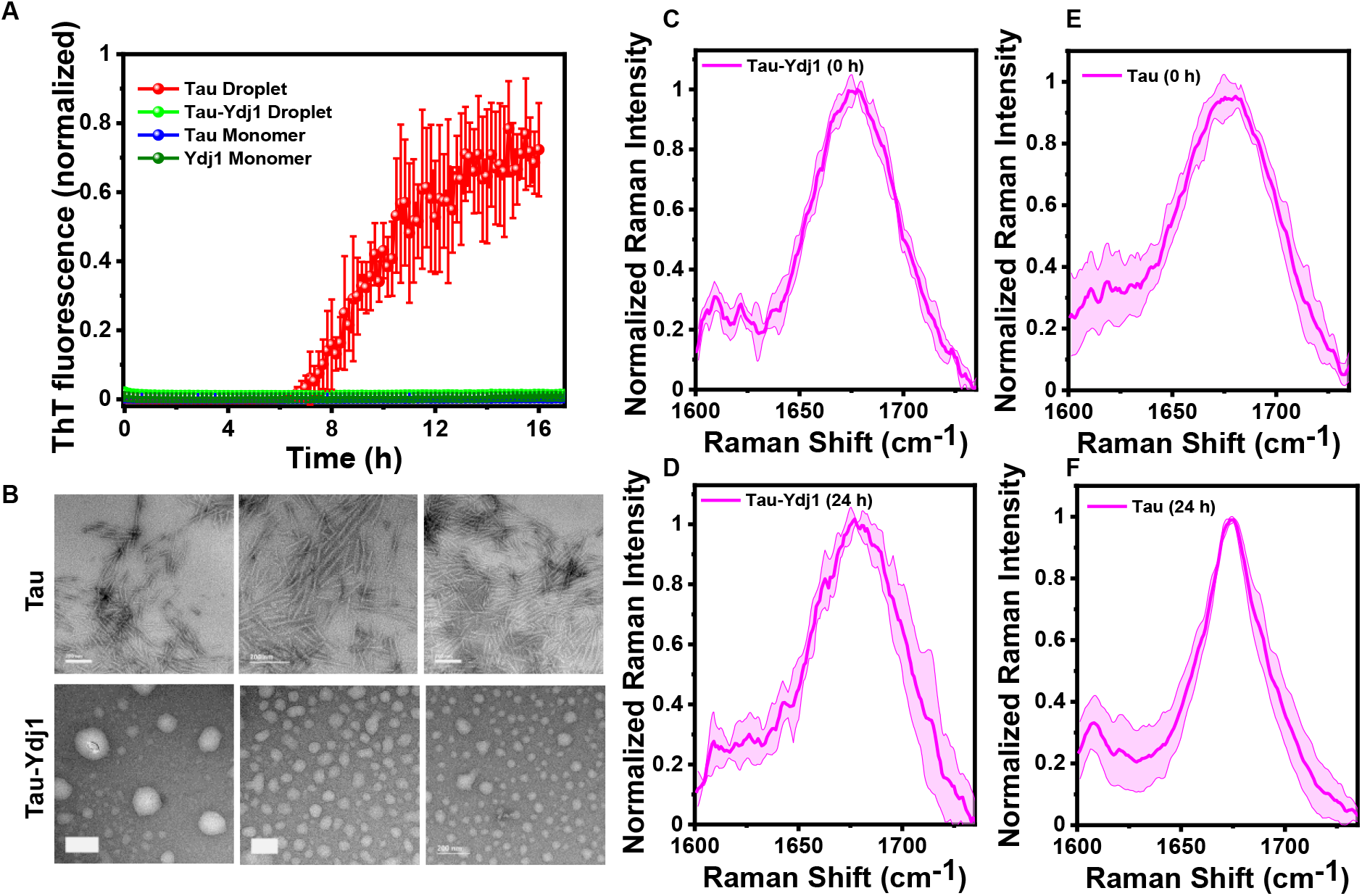
Ydj1 halts tau fibril formation. (*A*) ThT kinetics of phase separation-mediated aggregation of tau via a liquid-to-solid transition; and separately, tau-Ydj1 droplets, monomeric Ydj1, and monomeric tau. Wherever used, tau and Ydj1 concentrations were both equal to 10 μM. The data represent mean ± SD; n = 3 independent experiments. (*B*) TEM images of final species formed at the end of the aggregation kinetics of tau (*Upper*) and tau-Ydj1 condensates (*Lower*), indicating the presence of amyloid-like fibrils and proteinaceous oligomeric species in the absence and presence of Ydj1, respectively. The experiment was performed thrice with similar observations. (*C-F*) Amide I vibrational Raman band of the dense phase of tau-Ydj1 and tau recorded after 5 minutes of incubation and after 24 hours of incubation.

### Ydj1 induces conformational expansion in tau promoting their co-condensation

Monomeric tau is highly soluble and exists as a natively unstructured polypeptide. Previous studies showed that tau acquires globally compact “S-shaped” or “paper-clip“-like structures, in which both of its termini project onto the central MTBR region (77–79). However, tau is known to exhibit conformational expansion into “open” forms in the presence of cofactors, mutations, or during phase separation (80–82). Such expanded conformers are devoid of long-range intramolecular contacts and can expose the central hydrophobic region that is enriched in several amyloidogenic motifs allowing tau to undergo phase separation and aggregation. In order to directly observe the conformational changes of tau during its co-condensation with Ydj1, we deployed the single-molecule FRET methodology that offers a powerful tool to capture the conformational distribution, dynamics, and subpopulations (81–84). To install FRET pairs, we created two dual-cysteine tau constructs on the null-cysteine variant of full-length tau by introducing cysteine residues via site-directed mutagenesis in the MTBR region containing the central hydrophobic segments. These two FRET constructs with different intramolecular distances, Q244C-S322C (79 residues) and Q244C-S400C (157 residues), encompassed the R1-R3 and the R1-R′ regions, respectively (Fig. 6*A*). For our experiments, we stochastically labeled these double cysteine variants of tau with thiol-active donor and acceptor dyes namely, Alexa Fluor 488 maleimide (donor) and Alexa Fluor 594 maleimide (acceptor). Our single-molecule FRET results for the Q244C-S322C monomer exhibited an unimodal distribution with a mean FRET efficiency of ∼ 0.4 corresponding to a mean distance of ∼ 58 Å that is expected for 79-residue-separation for an expanded coil in a good solvent. However, the second pair (Q244C-S400C), encompassing the tau pseudo repeat in the R1-R′ region, exhibited a mean FRET efficiency of ∼ 0.27 corresponding to a mean inter-dye distance of ∼ 64 Å. This distance is shorter than that is expected for an expanded coil indicating a significant chain collapse due to the presence of R4 and R′ repeats (Fig. 6*B*, *C*). This observation is further corroborated by the constructed charge-hydrophobicity plot for the same construct suggesting that the R1-R′ region of tau is collapsed, possibly due to the potential electrostatic interactions between the MTBR and the C-terminal part of the repeat region (Fig. S6*A,* S6*B*).

**Fig. 6.**
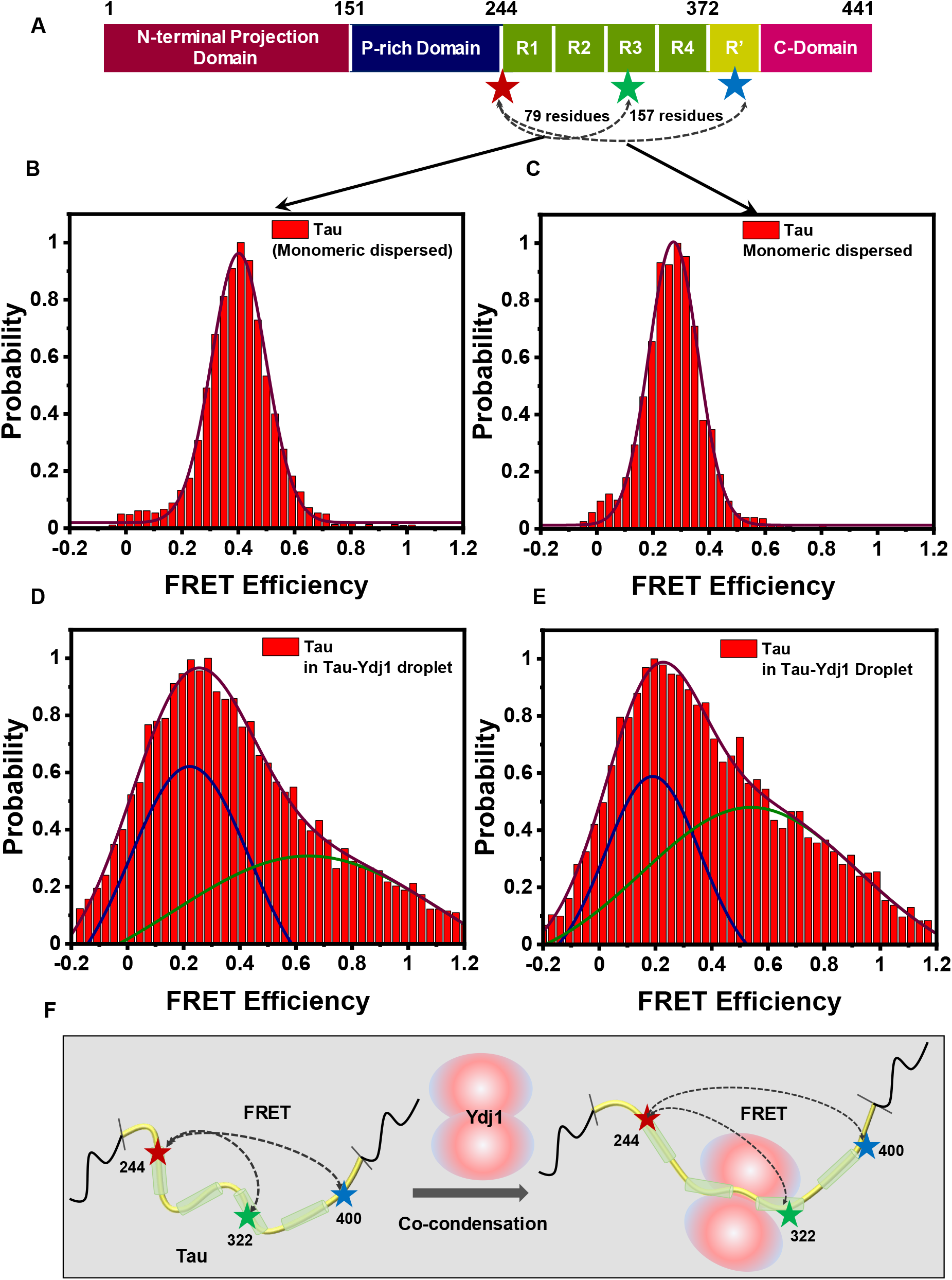
Single-molecule FRET dissects the conformational shape-shifting of tau in tau-Ydj1 condensates. (*A*) A schematic of the full-length tau constructs used for single molecule studies. Single-molecule FRET histograms of monomeric tau in the monomeric dispersed phase for dual-cysteine labeled constructs (*B*) Tau 244C-322C (inter-residue distance = 79 amino acids) and (*C*) Tau 244C-400C (inter-residue distance = 157 amino acids). Upon phase separation in the presence of Ydj1, tau undergoes conformational expansion, as seen from the broader histograms of tau in the tau-Ydj1 condensed phase for dual-labeled constructs (*E*) tau 244C-322C and (*F*) tau 244C-400C. The total number of events for all measurements was > 20,000, and the number of events at maxima was more than 700 in all cases. The binning time was 0.5 ms. (*G*) Schematic summarizing Ydj1 mediated conformational expansion in tau polypeptide chain.

Having established the monomeric conformations of tau, we next set out to capture the phase separation-induced conformational changes of tau using single-droplet single-molecule FRET experiments. Our measurements revealed that the tau polypeptide chain in the R1-R3 and the pseudo-repeat region (R′) underwent conformational expansion upon co-condensation into tau-Ydj1 droplets. While the conformational shift was significant for the R1-R3 region (244-322 pair) with a mean FRET efficiency of ∼ 0.2 (droplet) from ∼ 0.4 (monomeric dispersed), a very marginal shift was observed for the 244-400 pair with a FRET efficiency of ∼ 0.2 (droplet) from ∼ 0.3 (monomeric dispersed) (Fig. 6*D*, *E*). Such chain expansion events are known to offer more multivalent intermolecular contacts that are crucial for phase separation (83). We would like to note that both constructs exhibited a bimodal FRET efficiency distribution, with a smaller high-FRET population suggesting local structuring in the presence of Ydj1 with a FRET efficiency of ∼ 0.6 and ∼ 0.5 corresponding to 244-322 and 244-400, respectively. Such conformational heterogeneity possibly results from radial conformational preferences arising due to a “small-world” percolated network within biomolecular condensates having expanded and collapsed conformers (85). Additionally, conformational expansion in the R1-R3 region further supports our observation that the tau hexapeptide (PHF6) is essential for tau-Ydj1 interactions by accommodating Ydj1. Moreover, this region appears to have a higher conformational flexibility, as the 244-400 pair did not show any significant alteration in the FRET efficiency in the presence of Ydj1. A similar structural remodeling of tau has also been observed following its binding to tubulin and other components of the chaperone machinery, where although the overall dimensions of the tau repeat region remained unchanged, the lengths between individual repeats widened (76, 80, 82). Our results also indicated that polypeptide chains of tau possess higher conformational heterogeneity upon condensation, as indicated by a broader histogram, compared to a much sharper FRET histogram in the monomeric dispersed phase. The width of the histogram in the presence of Ydj1 can also be attributed to the fuzzy or dynamic nature of the tau-Ydj1 complex similar to other tau-chaperone complexes (75, 76). We note here that the width of FRET histograms in the condensed phase can also have some contributions from the photon shot noise and the orientation factor. Overall, these findings revealed that upon co-phase separation with Ydj1, tau undergoes conformational unwinding, enabling it to make a larger number of intermolecular contacts resulting in phase separation (Fig. 6*F*). Typically, upon tau phase separation, the enrichment of intermolecular interactions upon phase separation may lead to aggregation. However, upon the addition of Ydj1, the aggregation-prone hotspots of tau localized in the repeat region are occupied resulting in the inhibition of aberrant phase transition and aggregation into amyloid-like fibrils.

## Discussion

In this work, we showed the Hsp40-mediated regulation of phase separation and aggregation of tau. In the presence of tau, Ydj1 undergoes co-phase separation, leading to the emergence of highly dynamic, liquid-like condensates. The sequestration of Ydj1 within these condensates is associated with slower diffusion suggesting the formation of a densely crosslinked two-component protein network associated with partitioning this molecular chaperone (Fig. 1). Our findings indicated that tau-Ydj1 co-condensation is driven by both electrostatic and hydrophobic interactions that are corroborated by our observations on the modulation phase separation using phosphomimetic tau variants and other deletion constructs (Fig. 2 and 3). Our steady-state and picosecond time-resolved fluorescence anisotropy measurements captured the local ordering in the central MTBR of tau as indicated by lower anisotropy values for the N- and C-terminal ends of the tau chain in comparison to the central region. Our results also showed that RNA can alter the physical properties of tau-Ydj1 droplets by forming ternary condensates (Fig. 4). Additionally, our aggregation experiments demonstrated the Ydj1-mediated abrogation of tau aggregation and the transition of tau-Ydj1 liquid-like condensates into microscopic protein heterocomplexes (Fig. 5). Our study provides the mechanistic underpinning of chaperone-mediated regulation of aberrant phase transitions (Fig. 7).

**Fig. 7.**
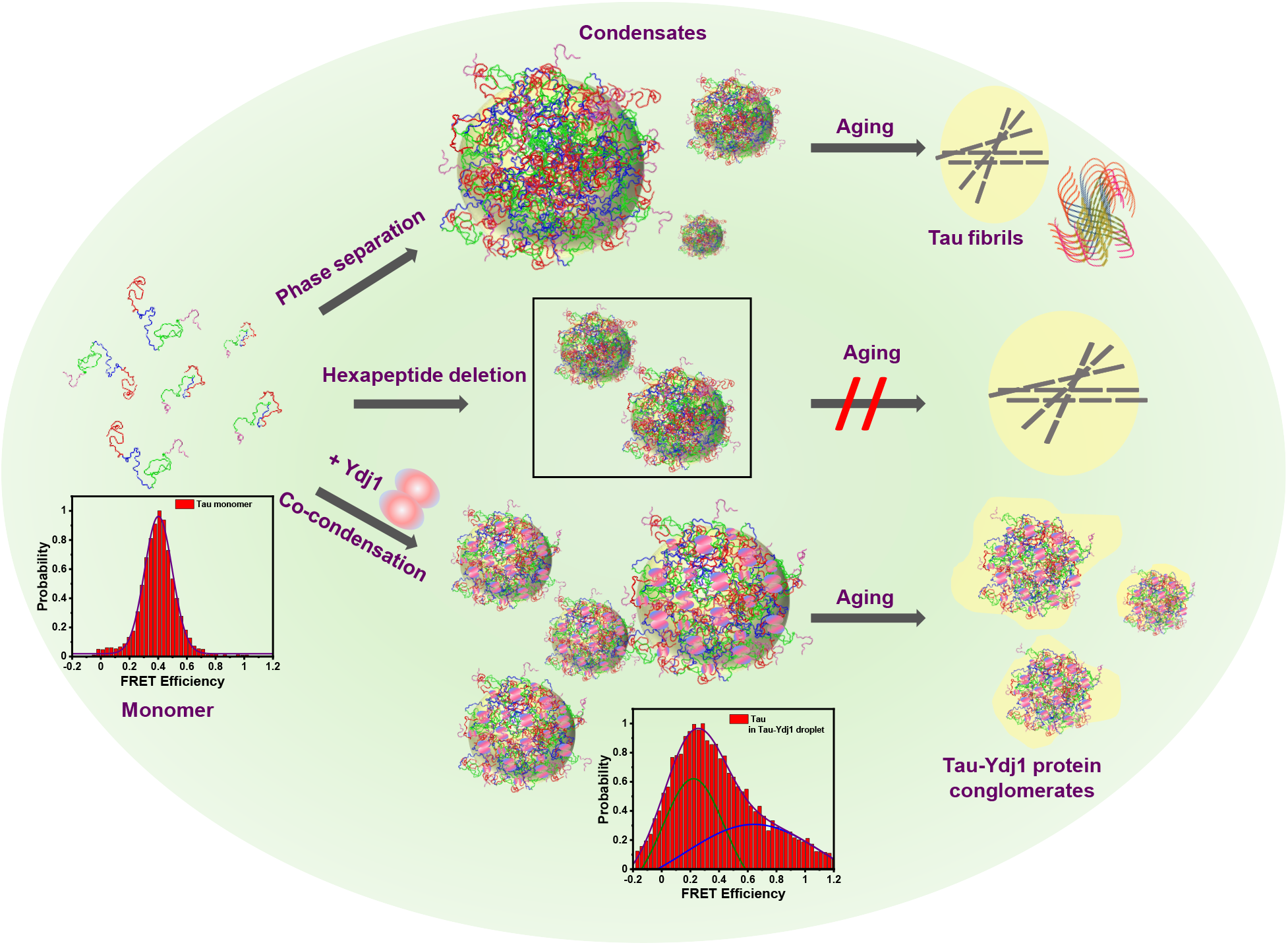
Schematic illustration of the regulation of tau phase transitions in the presence of Ydj1. Liquid-like phase-separated tau condensates can mature into solid amyloid-like aggregates implicated in neurodegeneration. This phase separation is associated with chain expansion and the exposure of the central MTBR region, containing hexapeptide amyloid motifs, which, in addition to regulation of aggregation, are also crucial for phase separation. Ydj1 modulates tau phase behavior by binding the exposed tau MTBR via its CTD and halting condensate maturation at an oligomeric stage.

The tau protein sequence incorporates well-characterized locally shielded and structured hexapeptide motifs, important for its phase separation and aggregation, out of which two lie at the interface of the R1/R2 and, separately, the R2/R3 regions of the MTBR (33, 79, 86). Any external stimuli, including mutations in the protein sequence, the presence of cofactors, or phase separation, are coupled to local structural re-ordering, promoting tau to adopt a pro-aggregation conformation. Separately, in our study, using selective hexapeptide deletion variants of tau, i.e., ΔPHF6* and ΔPHF6, we define the significance of these motifs for tau phase separation, both in the absence and presence of Ydj1, including the regulation of its phase transition into solid aggregates (Fig. S7). We showed that under our conditions, while ΔPHF6* had a phase behavior similar to full-length tau, the propensity of ΔPHF6 to undergo phase separation was lower, with droplet formation seen only at higher protein concentrations (Fig. S7*A*, *B*). Our saturation concentration measurement further validated this observation. We observed a higher C_sat_ for ΔPHF6 phase separation, underscoring the importance of the PHF6 region in driving tau condensation. Interestingly, while monitoring the thioflavin-T intensity during our aging experiments using these mutants, we noted the inhibition of aggregation for both tauΔPHF6* and ΔPHF6, with both variants showing similarly low ThT intensities in comparison to full-length phase-separated tau (Fig. S7*C*). These results indicate the importance of the cooperativity between the aggregation-promoting hexapeptides, with either deletion inhibiting aggregation. This observation, however, does not hold true for the initiation of tau phase separation. As inferred from our results, the PHF6 region of tau, which usually makes up the N-terminal region of tau amyloid cores, is critical for tau condensation. We hypothesize that the PHF6 sequence serves as a hotspot not only for governing tau aggregation as conventionally established but also on account of its significance in potentiating tau phase separation. This region has the highest propensity to form transient, beta-rich structured elements in the tau chain, potentially serving as sites for nucleating tau phase separation. On the contrary, for tau aggregation, the modular nature of these motifs in structuring tau warrants their cooperativity. This suggests that both amyloid-prone motifs must conjointly undergo a structural reconfiguration that is inhibited upon the deletion of either hexapeptide, allowing tau to retain its shielded anti-aggregation conformation. Moreover, in the context of the molecular chaperone-mediated shapeshifting of tau and its heterotypic phase separation with chaperones, including Hsps, the PHF6 region acts as a focal point of contact, as suggested by the diminution of tauΔPHF6 phase separation even in the presence of Ydj1 (Fig. 3).

Our single-molecule FRET studies captured conformational shapeshifting events experienced by the tau chain as it undergoes condensation in the presence of Ydj1 (Fig. 6). In solution, tau typically exists in a collapsed paperclip/S-shaped conformation, where the N and C-terminals of tau fold back upon the central MTBR region, consequently shielding the central hydrophobic region (77,78). Previously reported single-molecule studies on biomolecular condensates have reported the phase separation-associated conformational expansion of polypeptides, including tau (81, 83, 84). Based on our observations using the PHF deletion mutants of tau, we hypothesize that phase separation leads to the exposure of its central, highly hydrophobic, aggregation-prone repeat region. The resulting long-range inter-chain contacts now established, allow the creation of a dense viscoelastic network within the condensed phase. Over time, this network, with dominating contributions from the contacts promoted within the repeat region, can mature into a gel and, finally, a solid-like state, potentially generating beta-rich amyloid-like fibrils. Conversely, upon tau-Ydj1 coacervation, Ydj1 competes for the already-exposed hexapeptide region of tau lying in the MTBR with neighboring tau molecules. The sequestration of this region by Ydj1 makes it inaccessible for inter-chain interactions with neighboring tau, subsequently halting tau fibrilization. We propose that the highly dynamic and fuzzy nature of the tau-Ydj1 complex results in frequent association and dissociation events within the condensate, allowing tau molecules to establish prolonged contacts, leading to the formation of heterotypic proteinaceous “conglomerates” (Fig. 7).

In summary, our results highlight the self-sufficiency of the Hsp40 family of chaperones in abrogating tau maturation into fibrils. Cellular chaperoning is a complex process where multiple closely associated chaperones work in a feedback loop leading to clearance of toxic species and pathological aggregates (44–53). While complexes of major chaperone proteins, such as the Hsp70/Hsp90 complex, have been proven to be central in regulating tau aggregation and stability (46, 47), the role of the ATP-independent class of co-chaperones, such as Hsp40, in regulating the tau maturation via phase separation remains less understood. Our findings elucidate how molecular chaperones, as part of the cellular protein quality control mechanism, govern the conformational changes of phase-separated tau. Overall, our study underscores the broader significance of molecular chaperones in governing the properties of biomolecular condensates associated with physiology and disease.

## Materials and Methods

Detailed materials and methods are included in *Supporting Information*. The truncations of the tau protein and Ydj1 were created using full-length variants of the respective constructs. All mutations, including the single, double-cysteine, and deletion mutants of tau, were created using a QuikChange site-directed mutagenesis kit (Stratagene). The phosphomimetic mutants of tau were synthesized by Gene to protein. Tau protein was purified using a cation-exchange column followed by gel filtration, whereas cleavable N-terminal His-tagged Ydj1 was purified using a Co-NTA column, and His-tag was removed using TEV protease. Phase separation assays were performed using turbidity measurements and confocal microscopy. FRAP experiments were performed using ∼ 1% Alexa Fluor 488-C5-maleimide-labeled protein on a Zeiss LSM 980 super-resolution microscope coupled with an Elyra 7. Single-droplet anisotropy measurements and FCS experiments were performed on the MT200 time-resolved fluorescence confocal microscope (PicoQuant) using F5M and Alexa Fluor 488-C5-maleimide-labeled tau protein for the anisotropy and FCS measurements, respectively. The single-molecule FRET experiments using dual-cysteine tau constructs labeled with Alexa Fluor 488 and Alexa Fluor 594 as the FRET pair were also performed on MT200. Both data acquisition and analysis were performed on commercially available software, SymphoTime64 version 2.7 (PicoQuant). Aggregation kinetics were recorded on a POLARstar Omega Plate Reader Spectrophotometer (BMG Labtech) in NUNC 96-well plates. Vibrational Raman spectroscopic studies were performed on an inVia laser Raman microscope (Renishaw) using a 100× objective lens (Nikon) and a 785-nm near-infrared laser. TEM images were acquired following the negative of the sample on a Jeol JEM F-200 multi-purpose electron microscope.

## Acknowledgments

We thank IISER Mohali, Science and Engineering Research Board (SUPRA SPR/2020/000333, CRG/2021/002314, and J.C. Bose Fellowship JCB/2023/000016 to S.M.), Department of Science and Technology, Govt. of India (FIST grant # SR/FST/LS-II/2017/97 to the Department of Biological Sciences, IISER Mohali), Indo-French Centre for the Promotion of Advanced Research (IFC/A/6903-3/2023/680 to S.M.), and Ministry of Education, Govt. of India (Centre of Excellence grant to S.M.), Council of Scientific and Industrial Research (fellowship to S.K.R.), and the Prime Minister’s Research Fellowship to R.K. for financial support. Prof. Elizabeth Rhoades (University of Pennsylvania, USA) and Prof. Deepak Sharma (Institute of Microbial Technology, Chandigarh) for the kind gift of the DNA plasmids for full-length tau and Ydj1, respectively, Prof. N. Periasamy (Retd. TIFR Mumbai) for providing us with the fluorescence decay analysis program, Dr. Dipankar Bhowmik for helping us with schematic, and the members of the Mukhopadhyay lab for critically reading this manuscript.

## Authors Contributions

S.K.R. and S.M. conceived the project. S.K.R., R.K., A.S., and A.J. performed the experiments and analyses. S.K.R. prepared the figures. S.K.R. and R.K. wrote the first draft. S.M. supervised the work, edited the manuscript, obtained funding, and provided the overall direction. All authors discussed the results and commented on the manuscript.

