## Supplementary Information for "Chaperone-mediated heterotypic phase separation regulates liquid-to-solid phase transitions into amyloid fibrils"

### Materials and Methods

### Materials

2-(N-morpholino)ethanesulfonic acid (MES), glacial acetic acid, ammonium sulfate, 2-[4-(2-hydroxyethyl)piperazin-1-yl]ethanesulfonic acid (HEPES), sodium phosphate monobasic dihydrate, sodium phosphate dibasic dihydrate, tris base, sodium hydroxide, sodium chloride, magnesium chloride, potassium chloride, imidazole, Thioflavin T, 1,4-dithiothreitol (DTT), Triton X-100, PolyU sodium salt, phenylmethylsulfonyl fluoride (PMSF), TCEP (tris(2-carboxyethyl)phosphine), 1,6-Hexanediol, ethylenediaminetetraacetic acid (EDTA), nuclease-free water (NFW), were of highest purity grade, obtained from Sigma (St. Louis, MO, USA). Guanidinium hydrochloride was purchased from Amresco. Ampicillin, chloramphenicol, streptomycin sulfate, and isopropyl-β-thiogalactopyranoside (IPTG) were purchased from Gold Biocom (USA). All the fluorescent probes used in this study, mainly fluorescein-5-maleimide (F5M), Alexa Fluor 488-C5-maleimide, Alexa Fluor 488-NHS Ester (Succinimidyl Ester), Alexa Fluor 594-C5-maleimide, and Alexa Fluor 647-C2-maleimide, were obtained from Molecular Probes, Invitrogen. SP-sepharose, Q-sepharose, and Co-NTA resins were purchased from Qiagen. HiLoadTM Superdex-G75 16/600 prep grade (pg) column was obtained from GE Healthcare Life Sciences (USA). Amicon membrane filters for concentrating protein were purchased from Merck Millipore. All the buffer solutions were freshly prepared in Milli-Q water and filtered before use. The pH of each buffer solution was adjusted (± 0.02) at 25 °C using a Metrohm 827 lab pH meter.

#### Methods

##### Bioinformatic Analysis

The distribution of charges throughout both protein chains was analyzed using the Classification of Intrinsically Disordered Ensemble Regions (http://pappulab.wustl.edu/CIDER/analysis) CIDER (1) tool.

##### Site-directed mutagenesis and construct details

The constructs used in our experiments were created using the 6X His-Tau 2N4R-17C plasmid, which was a kind gift to us from Prof. Elizabeth Rhoades (University of Pennsylvania, USA). The 6xHis tag was removed from this construct via cloning, and all the variants of tau used in this study, including the single and double cysteine variants, the PHF6 and PHF6* deletion mutants, and the domain-specific mutants of tau (Nh2-tau (26-230) and tau truncation (151-399)) were created using this cloned construct via site-directed mutagenesis using a QuickChange kit (Stratagene). Additionally, unless mentioned otherwise, a null-cysteine variant of tau (C291S, C322S) was used for all our experiments. The phosphomimetic variants of tau (tau 17E and tau MARK) were synthesized using Gene to Protein. All the mutations were verified using sequencing. The yeast Ydj1 plasmid was a kind gift from Prof. Deepak Sharma (Institute of Microbial Technology, Chandigarh, India). The N (residues 1-207) and C-terminal domains (residues 208-409) of Ydj1 were cloned in the same vector using wild-type as a template. All the mutations were verified using sequencing.

##### Recombinant protein expression and purification

All tau variants were expressed in *E. coli* BL21(DE3) standard cells using 0.5 mM IPTG at 37°C for one hour. The proteins were purified in a native condition using ion exchange chromatography followed by size exclusion chromatography. Briefly, after expression, the cells were lysed via sonication followed by boiling in a lysis buffer containing 20 mM MES, 500 mM NaCl, 1 mM EDTA, 2 mM DTT, 1 mM $MgCl_2$, 1 mM PMSF, pH 6.5. The cell debris was removed using high-speed centrifugation, and the supernatant was treated with streptomycin sulfate and glacial acetic acid to remove nucleic acid contamination. The collected supernatant was incubated with 60% ammonium sulfate to precipitate the protein, which was isolated using high-speed centrifugation. The dried protein pellets were resuspended in Buffer A (20 mM MES, 50 mM NaCl, 1 mM EDTA, 2 mM DTT, 1 mM $MgCl_2$, 1 mM PMSF, pH 6.5) and loaded onto an SP-Sepharose column. The bound protein was eluted using a linear salt gradient that reached a final concentration of 100% of buffer B (20 mM MES, 1M NaCl, 1 mM EDTA, 2 mM DTT, 1 mM $MgCl_2$, 1 mM PMSF, pH 6.5). Nh2hTau was purified similarly using anion-exchange chromatography with Q-sepharose resin. The peak fractions were pooled together and further purified using size exclusion chromatography in buffer C (25 mM HEPES, 50 mM NaCl, 2 mM DTT, pH 7.4). The protein-containing fractions were concentrated and stored at -80 °C till further use. In the case of tau 17E and tau MARK, slightly different ion exchange buffers were used. Tau 17E protein pellets were resuspended in 25 mM HEPES, 50 mM NaCl, 1 mM EDTA, 2 mM DTT, pH 7.4, and loaded onto a Q-Sepharose column. The protein was eluted in a linear gradient with a 100% final concentration of buffer B (25 mM HEPES, 50 mM NaCl, 1 mM EDTA, and 2 mM DTT, pH 7.4). For tau MARK, buffer A contained 20 mM MES, 50 mM NaCl, 1 mM EDTA, pH 6.5 and was bound to an SP-Sepharose column, while buffer B contained 20 mM MES, 1 M NaCl, 1 mM EDTA, pH 6.5. As described above, these mutants were further polished by size exclusion chromatography in the presence of 2 mM DTT. All double cysteine variants were directly eluted in 6M GdMCl, 10 mM HEPES, 50 mM NaCl, 300 μM TCEP, and pH 7.4 during size exclusion chromatography. The proteins were concentrated using 10kDa MWCO Amicon membrane filters and stored at -80 °C till further use.

N-terminal 6X His-tagged recombinant pPROEX-Htb-Ydj1, Ydj1 N-terminal, and Ydj1-C-terminal fragments from *S. cerevisiae* were overexpressed in Rosetta DE3 *E. coli* cells using 0.3 mM IPTG at 15 °C for 14 hours. Harvested cells were resuspended in chilled lysis buffer (25 mM HEPES, 500 mM NaCl, 20 mM $MgCl_2$, 20 mM KCl, pH 7.4) and incubated with lysozyme (2 mg/ml) at 4 °C followed by sonication. The cell lysate was cleared using high-speed centrifugation, and purification was carried out using Co-NTA chromatography with an imidazole gradient. The 6X-His tag was removed overnight at 4 °C in 25 mM HEPES, 500 mM NaCl, 20 mM KCl,10 mM MgCl2, 1 mM DTT, pH 7.4 using in-house purified recombinant Tobacco Etch virus (TEV) protease. His-tag removal was carried out by passing the cleaved Ydj1 through a Co-NTA column. The eluted protein was concentrated using a 10 kDa MWCO Amicon membrane filter and stored at -80 °C until further use. Before experiments, Ydj1 was freshly dialyzed into 20 mM HEPES, 2 mM DTT, pH 7.4 using 10kDa MWCO Amicon membrane filters. The purity of all the proteins was validated using SDS-

PAGE, and freshly purified proteins were used for all experiments to avoid freeze-thaw cycles. The concentrations of the proteins were estimated using $\varepsilon_{280}$ = 6400 $M^{-1}cm^{-1}$ for full-length tau and tau ΔPHF6*, $\varepsilon_{280}$ = 2560 $M^{-1}cm^{-1}$ for Nh2htau and tau truncation, $\varepsilon_{280}$ = 5120 $M^{-1}cm^{-1}$ for tau ΔPHF6, and $\varepsilon_{280}$ = 23,475 $M^{-1}cm^{-1}$ for Ydj1, $\varepsilon_{280}$ = 12090 $M^{-1}cm^{-1}$ for Ydj1 C-terminal fragment, $\varepsilon_{280}$ = 7680 $M^{-1}cm^{-1}$ for Ydj1 N-terminal fragment respectively.

**Fluorescence labeling**

All labeling reactions were carried out using thiol-reactive maleimide labeling. Single cysteine variants of tau were site-specifically labeled under denaturing conditions (6M GdMCl, 10 mM HEPES, 50 mM NaCl, pH 7.4) using fluorescein-5-maleimide (F5M) and Alexa Fluor C5-maleimide dyes (Alexa Fluor 488 and Alexa Fluor 594) for anisotropy and imaging experiments, respectively. For F5M labeling, proteins were mixed in a 10:1 ratio (dye: protein); for Alexa fluor labeling, this ratio was 2:1 (dye:protein). The reactions were incubated in the dark for 2-3 hours while stirring and were dialyzed into 25 mM HEPES, 50 mM NaCl, pH 7.4 using 10 kDa MWCO centrifugal filters.

For dual labeling of the double cysteine variants of tau, the denatured protein was incubated on a stirrer with the donor (Alexa Fluor 488) dye in a 0.8:1 ratio (dye:protein) for ~ 2 h at room temperature in the dark. The acceptor dye (Alexa Fluor 594) was added in a 0.8:5 ratio (donor: acceptor), and the reaction mixture was incubated overnight at 4 °C with stirring. The next day, the free dye was removed by buffer exchange of the protein (6M GdMCl, 10 mM HEPES, 50 mM NaCl, pH 7.4) using 10kDa MWCO centrifugal filters. Finally, the dual-labeled protein was buffer-exchanged into 25 mM HEPES, 50 mM NaCl, and 2 mM DTT, pH 7.4. Ydj1 was labeled non-specifically under native conditions using Alexa Fluor 647-C2-maleimide and Alexa Fluor 488-C5-maleimide (1:1) for imaging and FRAP experiments. The concentrations of the labeled proteins were measured using $\varepsilon_{495}$ = 68,000 $M^{-1}cm^{-1}$, for F5M, $\varepsilon_{495}$= 72,000 $M^{-1}cm^{-1}$, for Alexa Fluor 488-C5-maleimide, $\varepsilon_{590}$ = 92,000 $M^{-1}cm^{-1}$ for Alexa Fluor 594-C5-maleimide, and $\varepsilon_{647}$ = 2,39,000 $M^{-1}cm^{-1}$ for Alexa Fluor 647-C2-maleimide. For polyU RNA labeling, the 5'-end of RNA was activated using NHS-EDC coupling in pH 6.5 buffer followed by buffer exchange in MiliQ water using a 10 kDa MWCO Amicon membrane filter. The labeling of RNA was done by diluting the final reaction mixture in a buffer (20 mM Sodium phosphate, 50 mM NaCl, pH 7.4) prepared using MiliQ water. A two-fold molar excess of Alexa Fluor488 NHS Ester (Succinimidyl Ester) was used for labeling at room temperature. Excess free dye was removed using a membrane filter. Labeled RNA concentration was estimated by monitoring absorbance at 260 and 495 nm (absorbance maxima of fluorophore).

**Phase separation assays**

The concentration of the stock of tau was kept constant at 360 μM throughout the experiments. Tau phase separation was induced in the reaction buffer (20 mM HEPES, 2 mM DTT, pH 7.4) by diluting the stock to 10μM. For tau-Ydj1 condensates, 10 μM tau was added to Ydj1 diluted to 10 μM in the same buffer. Phase separation was quantified by measuring the turbidity of the reactions at 350 nm, 25 °C on a MultiskanGo (Thermo scientific) plate reader using 96-well NUNC optical bottom plates. The sample volume was kept constant at 100 μL. For 1,6-Hexanediol dependent turbidity measurements, the required concentration of hexanediol was introduced into the reaction mixture from a 50 % (weight/volume) stock solution, and data were acquired as described above. For RNA-dependent turbidity measurements and with truncation variants, Nanodrop (Genova Life Science Spectrophotometer, ver 1.51.4) was used for the same. Data was plotted without any background subtractions. The mean and standard deviations were obtained from three independent sets of measurements recorded on the same day.

**Saturation concentration ($C_{sat}$) estimation using sedimentation assays**

The tau droplet and tau-Ydj1 reactions were set up and incubated at room temperature for 10 minutes, followed by high-speed centrifugation at 16,400 rpm at 25 °C for 35 minutes. The reactions were doped with 10% of F5M labeled tau. Following centrifugation, the supernatant was collected and diluted with a salt-containing buffer (25 mM HEPES, 50 mM NaCl, 2 mM DTT, pH 7.4) to prevent the induction of phase separation. Protein concentration was estimated in the light phase by monitoring absorbance at 495 nm. The suitable dilution factor and a factor of 10 were incorporated into the calculations to obtain the Csat (2).

**Confocal microscopy imaging**

All the reactions were imaged at room temperature using a ZEISS LSM 980 Elyra 7 super-resolution microscope equipped with a high-resolution monochrome cooled AxioCamMRm Rev. 3 FireWire(D) camera, using a 63x oil-immersion objective (numerical aperture 1.4). All Airyscan confocal imaging experiments were performed using the associated Airyscan 2 detector (32 channels GaAsP). For droplet reactions, the unlabeled proteins were doped with 1% labeled proteins in the reaction buffer and imaged using a 488 nm laser diode (11.9 mW) and a 590 nm excitation source for Alexa fluor 488 and Alexa fluor 594, respectively. For three-color imaging, a 632 nm excitation source for Alexa fluor 647, in addition to two other excitation sources, were used. The images were acquired using 1840 × 1840 pixels resolution at a 16-bit depth. The obtained images were processed and analyzed using the Zen Blue 3.2 (3.2) software.

**Fluorescence recovery after photobleaching (FRAP)**

FRAP experiments were carried out using the confocal imaging setup described above. For all such experiments, unlabeled proteins were doped with the corresponding 1% of Alexa Fluor 488 labeled proteins, and a region of 1 μm diameter was bleached using the 488 laser diode. The recovery was recorded using the ZEN blue 3.2 (ZEISS) software, and the obtained recovery profiles were background corrected, normalized, and plotted using Origin 2020b. At least 3 independent sets of experiments were performed, and 5-7 traces were used for creating the plots.

**Single-droplet steady-state and time-resolved anisotropy measurements**

Single-droplet anisotropy measurements were recorded using the PicoQuant MicroTime (MT200) microscope as described above. Monomeric tau and phase-separated tau-Ydj1 reactions were freshly prepared using F5M-labeled single cysteine variants of tau, which were excited using a 488 nm laser. The emitted fluorescence was selectively collected using a 520/35 bandpass filter, out-of-focus light was filtered out using a 50μM pinhole, and the collected light was divided by a polarizer into two detectors (SPADs). Anisotropy imaging was performed for steady-state measurements, while point time traces were collected for time-resolved information. Correction factors required for data analysis were calculated using measurements with free dye solutions. In the case of tau-Ydj1 reactions, droplets were chosen as regions of interest (ROI) using the associated SymphoTime64 v2.7 software. Data acquisition and analysis were performed on the same software. Steady-state anisotropy is given by

$$r_{ss} = \frac{I_{\perp} - I_{\parallel}}{[1-3L2]I_{\parallel} + [2-3L1]I_{\perp}} \quad \text{--- (1)}$$

where $I_{\parallel}$ and $I_{\perp}$ are the background corrected parallel and perpendicular fluorescence intensities, and L1 and L2 are the correction factors for the used objective lens.

The time-resolved anisotropy decay profiles that were acquired were fitted globally using the following relations:

$$I_{\parallel}(t) = 1/3I(t)[1 + 2r(t)] \quad \text{--- (2)}$$

$$I_{\perp}(t) = 1/3I(t)[1 - r(t)] \quad \text{--- (3)}$$

Here, $I$ denotes the time-dependent fluorescence intensity collected at the magic angle (54.7°) geometry.

The collected intensity decays could be approximated to a biexponential decay model as follows:

$$r(t) = r_0[\beta_1 e^{\left(\frac{-t}{\phi_1}\right)} + \beta_2 e^{\left(\frac{-t}{\phi_2}\right)}] \quad \text{--- (5)}$$

where $\phi_1$ and $\phi_2$ denote the fast and the slow rotational correlation times associated with the local and global dynamics of the associated protein chain, respectively. $\beta_1$ and $\beta_2$ denote the respective amplitudes associated with these fast and slow rotational correlation times, while $r_0$ denotes the intrinsic time-zero fundamental anisotropy of the fluorophore. Reduced $\chi^2$ values were used to assess the goodness of fit (3).

**Fluorescence correlation spectroscopy**

FCS measurements were performed on the abovementioned instrument (PicoQuant MT200). The confocal volume ($V_{eff}$) and the corresponding structural parameter ($\kappa$) for our system were determined using a 1 nM solution of Alexa488, which gave us $V_{eff} > 1$ fL and $\kappa = 4.7$. These parameters were used as calibration values while curve-fitting data for the monomers and droplets. Dispersed monomeric and droplet solutions (tau, Ydj1, and tau-Ydj1) were prepared by mixing nanomolar concentrations of the Alexa488 labeled single cysteine variants of respective proteins with unlabeled proteins. Freshly prepared reaction mixtures (50 μL) were spotted onto coverslips, and measurements were performed. In the case of monomer, experiments were performed 50 μm inside the solution, whereas individual droplets were focused in the case of phase-separated solutions. Correlation curves (G(t)) were fitted using the triplet model.

$$G(t) = \left[1 + T\left[e^{\left(\frac{-t}{\tau_{Trip}}\right)} - 1\right]\right] \sum_{i=0}^{n_{Diff}-1} \frac{\rho[i]}{\left[1+\frac{t}{\tau_{Diff}[i]}\right]\left[1+\frac{t}{\tau_{Diff}[i]\kappa^2}\right]^{0.5}} \quad \text{--- (6)}$$

where G(t) is the correlation amplitude, $\rho$ denotes the contribution of the $i^{th}$ diffusing species, T denotes the fraction of the triplet state, $\tau_{Trip}$ is the lifetime of the triplet state, $\tau_{Diff}$ is the diffusion time of the $i^{th}$ diffusing species, and $\kappa$ is the structure parameter of the corresponding confocal volume.

**Single-molecule FRET measurements and data analysis**

Single-molecule FRET experiments were performed on the Picoquant Microtime 200 (MT200) inverted time-resolved fluorescence confocal microscope. All experiments were performed in the PIE (Pulsed Interleaved Excitation) mode using pulsed laser sources (488 and 594) with a frequency of 20 MHz. The reactions were drop cast on 0.15 mm thick coverslips (#1) and placed on a Super Apochromat 60x water immersion objective with 1.2 NA. The emitted fluorescence was collected using the same objective, filtered using a 50μM pinhole, and subsequently separated into donor and acceptor channels using a dichroic mirror (zt594rdc). Optical filters were placed before the two SPADs (single avalanche photodiodes) that were used as detectors (530/50 BP for the donor and 645/75 BP for the acceptor). Dual-labeled tau was diluted to a final concentration of ~ 150 pM in 25 mM HEPES, 50 mM NaCl, 2 mM DTT, pH 7.4 buffer, and doped with 50 nM of unlabeled protein for monomer measurements to achieve surface passivation. For droplet reactions, 5-10 pM dual labeled protein was doped in a protein mixture containing 10 μM of both unlabeled tau and Ydj1 (4). Propyl gallate was used at a concentration of 250 μM in the reaction buffer as an oxygen scavenger to improve the photostability of the dye pair. A binning time of 0.5 ms and a minimum of 35 counts was used as a threshold for the obtained bursts for further analysis. FRET was calculated as:

$$E = \frac{I_A - \alpha}{\gamma I_D + (I_A - \alpha)} \quad \text{--- (7)}$$

Where $I_D$ and $I_A$ are the intensities in the donor and acceptor channels, respectively, and $\gamma$ accounts for the difference in the quantum yields and detection efficiencies of the donor and acceptor. $\alpha$ signifies the donor fluorescence leakage into the acceptor channel.

**Aggregation kinetics**

All Thioflavin T-based aggregation kinetics were recorded on a POLARstar Omega Plate Reader Spectrophotometer (BMG LABTECH, Germany) in NUNC-96 well plates. Each well was incubated with 150 μl of the reaction supplemented with a glass bead having a 3 mm diameter. ThT was used at a final concentration of 20 μM for all reactions. The kinetics were recorded under continuous stirring conditions at 100 rpm. The ThT intensity was plotted using Origin 2020b.

**Transmission Electron Microscopy**

The end product of the tau droplet and tau-Ydj1 droplet reactions set up for aggregation kinetics measurements were taken from the plate reader and pelleted down by centrifugation at 16,400 rpm, 25 °C. After pelleting down, the supernatant was discarded, and pellets were dissolved in 30 µl of reaction buffer. A small volume (5 µl) of the reactions was drop cast on 300-mesh carbon-coated electron microscopy grids and incubated for 5 minutes. Negative staining with 5 µl of uranyl acetate (1% w/v) was performed, followed by overnight incubation at room temperature after excessive stain removal. Imaging was performed on Jeol JEM F-200.

**Raman spectroscopy**

All Raman measurements were carried out using the dense phase from reaction mixtures. After incubation of the reaction mixtures (600 µL volume), these were subject to high-speed centrifugation at specific time points. The dense phase (pellet) was subsequently resuspended in 5 µL of 20 mM sodium phosphate buffer, pH 7.4, and deposited onto a glass slide covered with an aluminum sheet. Spectra were recorded for half-dried samples using an inVia laser Raman microscope (Renishaw, UK). Using a 100x objective lens (Nikon, Japan), the samples were focused and excited via a 785-nm NIR laser, with an exposure time of 10 s and 100% laser power. Spectra were recorded for tau and tau-Ydj1 droplets at different time points. Rayleigh scattering was filtered using an edge filter of 785 nm. The Raman scattering was collected and dispersed using a 1200 lines/mm diffraction grating and detected using an air-cooled CCD detector. The instrument's in-built Wire 3.4 software was used for data acquisition. All the data were averaged over 10 scans. Acquired spectra were baseline corrected and smoothened using Wire 3.4. Spectra were plotted using Origin 2020b (5).

**Table S1.** Primers used for creating point mutations and truncations:

| | |
|---|---|
| T17C Forward | GGAAGATCACGCTGGGACTTACGGGTTGGGGG |
| T17C Reverse | CCCCCAACCCGTAAGTCCCAGCGTGATCTTCC |
| S56C Forward | CACTGAGGACGGATGTGAGGAACCGGGC |
| S56C Reverse | GCCCGGTTCCTCACATCCGTCCTCAGTG |
| A158C Forward | CACCGCGGGGAGCATGCCCTCCAGGCCAG |
| A158C Reverse | CTGGCCTGGAGGGCATGCTCCCCGCGGTG |
| S199C Forward | CAGCGGCTACAGCTGCCCCGGCTCCCC |
| S199C Reverse | GGGGAGCCGGGGCAGCTGTAGCCGCTG |
| Tau ΔPHF6* Forward | CCAGCCGGGAGGCGGGAAGAAGCTGGATCTTAGCAACG |
| Tau ΔPHF6* Reverse | CGTTGCTAAGATCCAGCTTCTTCCCGCCTCCCGGCTGG |

| | |
|---|---|
| Tau ΔPHF6 Forward | CGTCCCGGGAGGCGGCAGTCCAGTTGACCTGAGC |
| Tau ΔPHF6 Reverse | GCTCAGGTCAACTGGACTGCCGCCTCCCGGGACG |
| Nh2-tau Forward | ATATATCATATGCAGGGGGGCTACACCATGCACC |
| Nh2-tau Reverse | ATATATCTCGAGTCAACGGACCACTGCCACCTTCTTGG |
| Tau (151-399) Forward | ATATATCATATGATCGCCACACCGCGGGG |
| Tau (151-399) Reverse | ATATATCTCGAGTCACTCCGCCCCGTGGTCTGTCTTGG |
| S291C Forward | GCAACGTCCAGTCCAAGTGCGGCTCAAAGG |
| S291C Reverse | CCTTTGAGCCGCACTTGGACTGGACGTTGC |
| S322C Forward | GAGCAAGGTGACCTCCAAGTGCGGCTCATTAGGC |
| S322C Reverse | GCCTAATGAGCCGCACTTGGAGGTCACCTTGCTC |
| S400C Forward | GTCGCCAGTGGTGTGTGGGGACACGTCTC |
| S400C Reverse | GGAGACGTGTCCCCACACACCACTGGCGAC |
| S433C Forward | GCTGACGAGGTGTGTGCCTCCCTGGCC |
| S433C Reverse | GGCCAGGGAGGCACACACCTCGTCAGC |
| CTD Forward | ATATATCTCGAGTCATTGAGATGCACATTGAACACC |
| CTD Reverse | ATATATCATATGAAAGTTGAAAACGAAAGGAAGATCC TAGAAGTCCATG |

**Table S2.** Recovered parameters from time-resolved fluorescence anisotropy decay analyses.

| **Tau-S244C-F5M** | **$\phi_1$ ($\beta_1$)** | **$\phi_2$ ($\beta_2$)** |
|---|---|---|
| Tau monomer | 0.89 ± 0.05 ns (0.59 ± 0.011) | 4.93 ± 0.39 ns (0.41± 0.015) |
| Tau-Ydj1 droplets | 0.99 ± 0.05 (0.32 ± 0.012) | 41.17 ± 1.24 (0.68 ± 0.013) |
| **Tau-S400C-F5M** | **$\phi_1$ ($\beta_1$)** | **$\phi_2$ ($\beta_2$)** |
| Tau monomer | 0.93 ± 0.03 ns (0.59 ± 0.011) | 5.03 ± 0.29 ns (0.41± 0.011) |
| Tau-Ydj1 droplets | 1.05 ± 0.04 (0.33 ± 0.011) | 39.87 ± 1.64 (0.67 ± 0.011) |

#### Supporting Information Figures

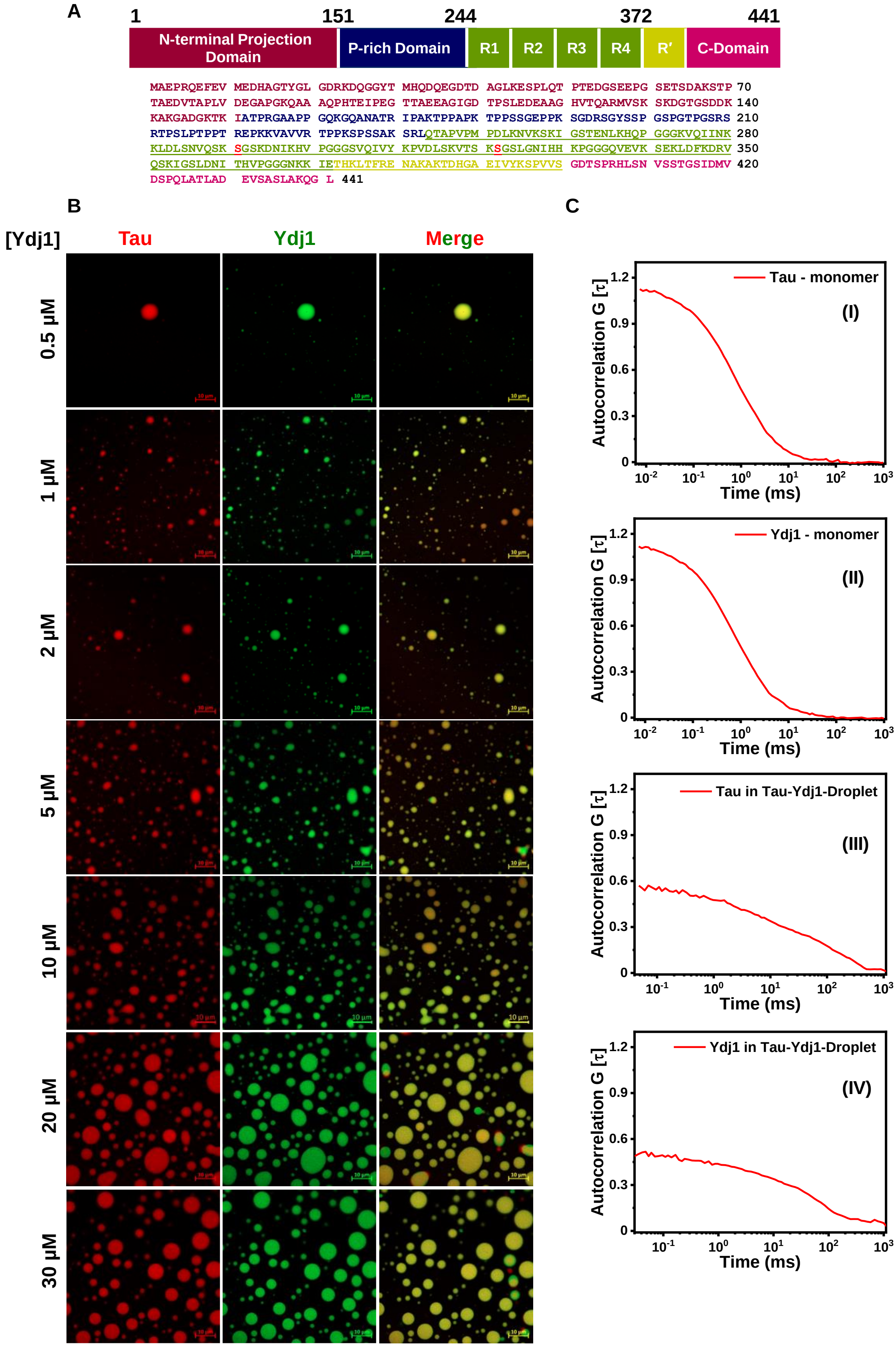


**Fig. S1. Tau phase separation is modulated by Ydj1.** (*A*) Domain architecture and amino acid sequence of tau. (*B*) Two-color Airyscan confocal images of tau (tau-Q244C-Alexa Fluor

594, red) and Ydj1 (sparsely labeled with Alexa Fluor 488, green) heterotypic condensates in the presence of an increasing concentration of Ydj1 (Scale bar, 10 µm). (*C*) Unnormalized FCS autocorrelation plots of tau monomer (I), Ydj1 monomer (II), tau in tau-Ydj1 droplets (III), and Ydj1 in tau-Ydj1 droplets (IV). The normalized versions of these plots are shown in Fig. 1*M*.

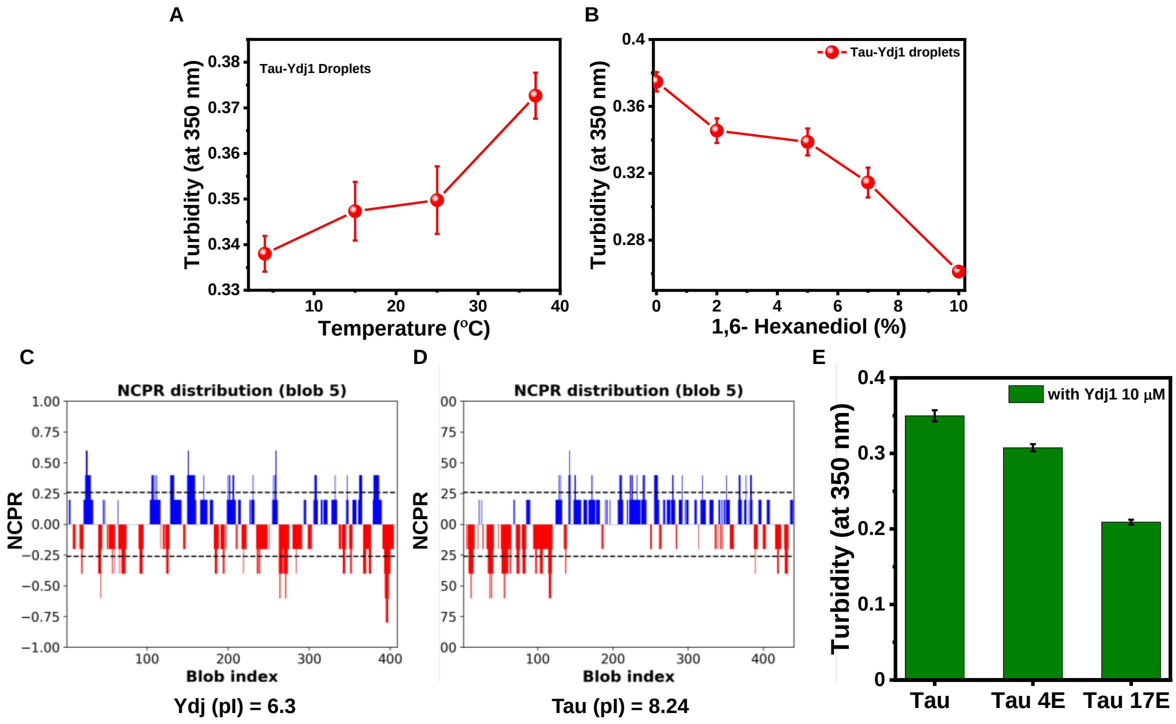


**Fig. S2. An interplay of electrostatic and hydrophobic interactions drives tau-Ydj1 condensation.** (*A*) Turbidity (O.D. at 350 nm) measurements of the tau-Ydj1 reaction mixture as a function of increasing temperature (Tau and Ydj1 concentrations were both kept fixed at 10 µM). The data represent mean ± SD; n = 3. (*B*) Turbidity (O.D. at 350 nm) measurements of the tau-Ydj1 reaction mixture as a function of increasing concentrations of 1,6-Hexanediol (Tau and Ydj1 concentrations were both kept fixed at 10 µM). The data represent mean ± SD; n = 3. (*C*) Net charge per residue profile of Ydj1 and (*D*) tau. (*E*) Turbidity values (O.D. at 350 nm) of full-length tau in comparison to tau-4E and tau-17E in the presence of 10 µM of Ydj1. The concentrations of all tau variants were kept fixed at 10 µM. The data represent mean ± SD; n = 3.

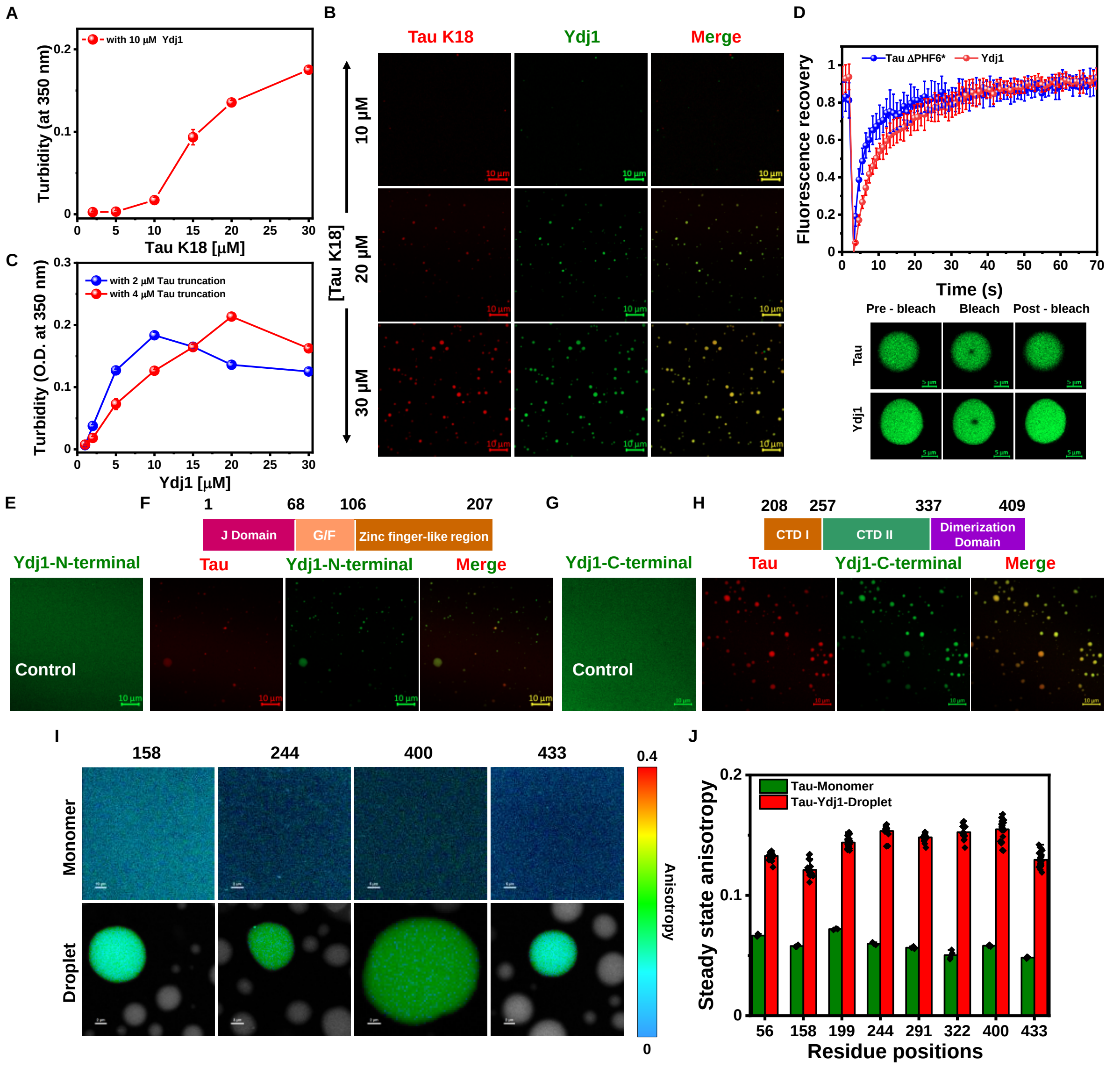


**Fig. S3. Domain-specific interactions are fundamental to tau-Ydj1 phase separation.** (*A*) Turbidity (O.D. at 350 nm) measurements of tau K18-Ydj1 reaction mixtures as a function of increasing concentration of Ydj1 (Tau K18 concentration was kept fixed at 10 μM). The data represent mean ± SD; n = 3. (*B*) Two-color Airyscan confocal images of tau K18 (tau-truncation A158C-Alexa Fluor 594, red) and Ydj1 (sparsely labeled with Alexa Fluor 488, green) complex coacervates in the presence of an increasing concentration of tau K18 (Scale bar, 10 μm). The concentration of Ydj1 was kept fixed at 10 μM. (*C*) Turbidity (O.D. at 350 nm) measurements of tau-truncation-Ydj1 reaction mixtures as a function of increasing concentration of Ydj1 (Tau truncation concentration was kept fixed at 2 μM (*blue*) and 4 μM (*red*). The data represent mean ± SD; n = 3. (*D*) FRAP kinetics of tau ΔPHF6*-Ydj1 droplets. Alexa Fluor 488-labeled proteins were used for both tau ΔPHF6* and Ydj1 for independent FRAP studies (Both tau ΔPHF6* and Ydj1 concentrations were 10 μM; tau-Q244C-Alexa Fluor 488 was used for tau ΔPHF6* bleaching). The data represent mean ± SD; n = 5. The corresponding fluorescent images acquired during FRAP measurements are shown below (Scale bar, 5 μm). (*E*) Airyscan confocal image of the Ydj1 N-terminal fragment (residues 1-207) (10 μM, green) (Scale bar, 10 μm). (*F)* Depiction of the Ydj1 N-terminal domain fragment and two-color Airyscan confocal images of tau (10 μM, red) and Ydj1-NTD (10 μM, green) (Scale bar, 10 μm). (*G*) Airyscan confocal image of the Ydj1 C-terminal fragment (residues 208-409) (10 μM, green) (Scale bar, 10 μm). (*H*) Depiction of the Ydj1 C-terminal domain (CTD) fragment and two-color Airyscan confocal images of tau (10 μM, red) and Ydj1-CTD (10 μM, green) (Scale bar, 10 μm). (*I*) Representative single-droplet steady-state fluorescence anisotropy images showing anisotropy heatmap of F5M labeled single-Cys mutants of tau spanning the sequence in the dispersed monomeric (Upper) and in tau-Ydj1 droplets (*Lower*). (*J*) Single-droplet steady-state fluorescence anisotropy measurements of F5M labeled single-Cys mutants of tau spanning the entire full-length sequence in the dispersed, monomeric state (olive) and in tau-Ydj1 droplets (red). Data for more than n = 30 different droplets were considered for droplet anisotropy.

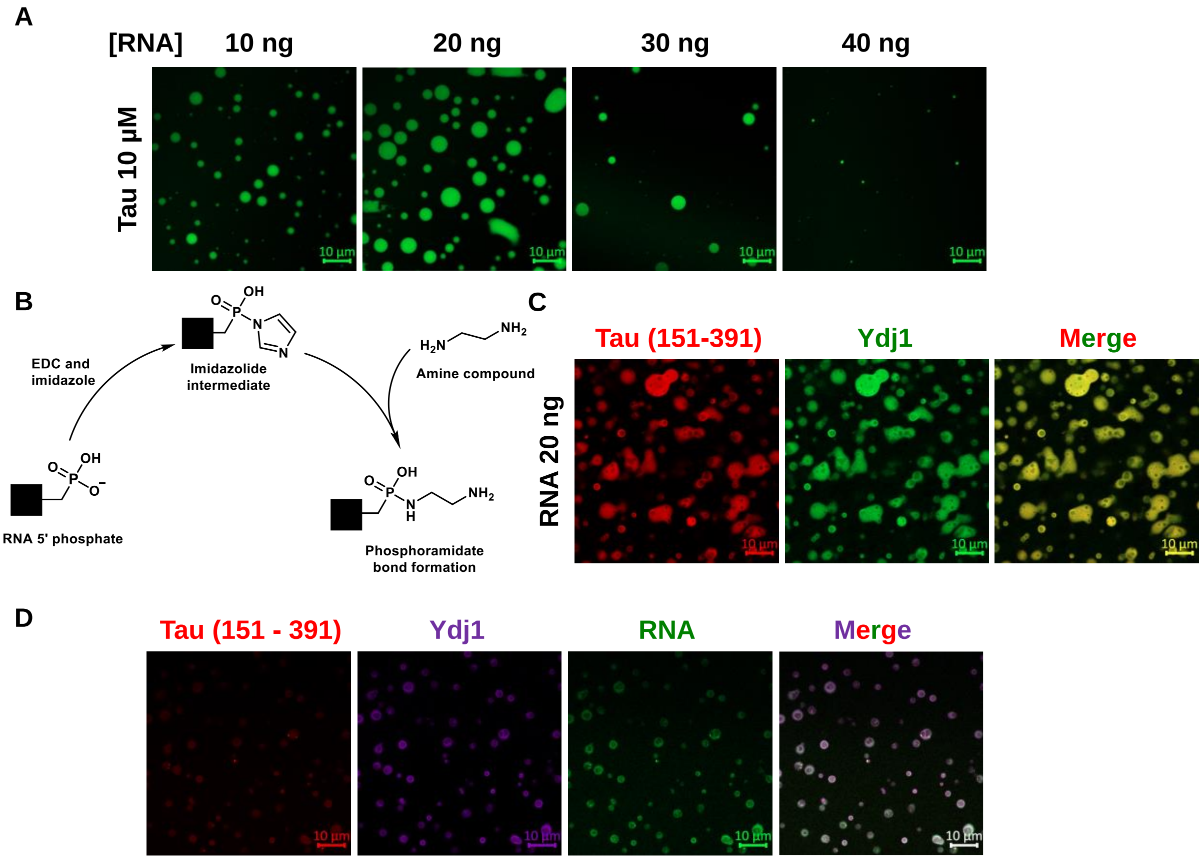


**Fig. S4. RNA-mediated tau-Ydj1 reentrant phase behavior.** (*A*) Airyscan confocal images of tau (tau-Q244C-Alexa Fluor 488, green) complex coacervates in the presence of an increasing concentration of polyU RNA (Scale bar, 10 µm). The concentration of tau was kept fixed at 10 µM. (*B*) Schematic depicting the labeling of RNA with the Alexa Fluor 488-NHS ester (succinimidyl ester) dye via the EDC-NHS-coupling mediated activation of its 5' phosphate (Image drawn using ChemDraw). (*C*) Two-color Airyscan confocal images of tau-truncation (tau-truncation A158C-Alexa Fluor 594, red) and Ydj1 (sparsely labeled with Alexa Fluor 488, green) complex coacervates in the presence of 20 ng/µL polyU RNA (Scale bar, 10 µm). The concentration of tau-truncation and Ydj1 was kept fixed at 10 µM. (*D*) Three-color Airyscan confocal image of colocalized tau-truncation (tau-truncation A158C-Alexa Fluor 594, red); Ydj1 (Alexa Fluor 647, pink) and RNA (5' Phosphate-Alexa Fluor 488, green) in tau-Ydj1-RNA coacervates (Scale bar, 10 µm). Both tau and Ydj1 concentrations were 10 µM, while RNA concentration was 20 ng/µL.

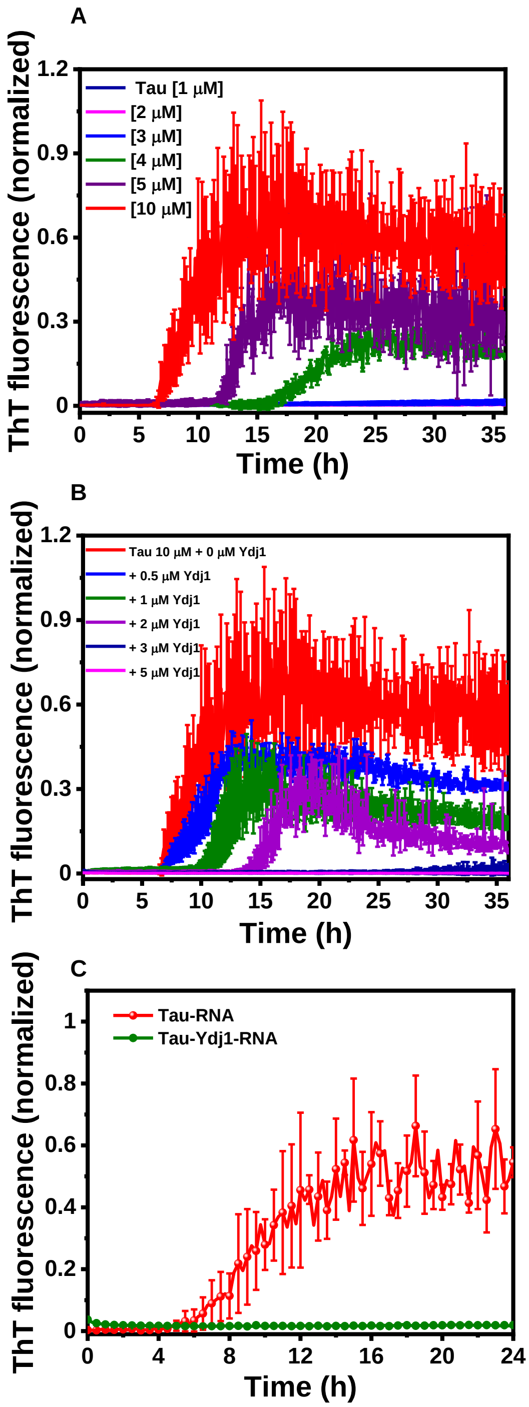


**Fig. S5. Ydj1 halts tau fibril formation.** (*A*) ThT kinetics of phase separation mediated aggregation of tau at varying concentrations. (*B*) ThT kinetics of the modulation of the liquid-to-solid transition of tau in the presence of varying concentrations of Ydj1. The concentration of tau was kept fixed at 10 μM. The same kinetics as in (A) have been used in the case of tau 10 μM for comparison. (*C*) ThT kinetics of phase separation-mediated aggregation of tau-RNA via a liquid-to-solid transition and, separately, tau-Ydj1-RNA droplets. Wherever used, tau and

Ydj1 concentrations were equal to 10 μM, and 20 ng/μL RNA concentration was used. All data represent mean ± SD for n = 3 independent experiments.

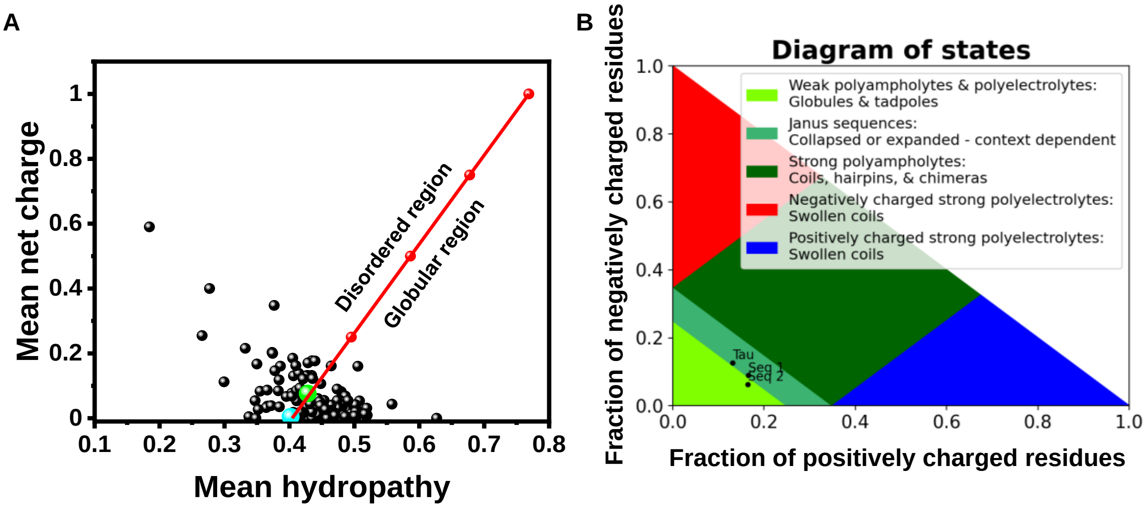


**Fig. S6. Single-molecule FRET dissects the conformational shape-shifting of tau in tau-Ydj1 condensates.** (*A*) Mean net charge as a function of mean hydrophobicity for a range of natively ordered and disordered proteins. The tau null cysteine and Q244C-S400C variants are represented in cyan and green, respectively. (*B*) A sequence annotated diagram of IDP states based on the fraction of negative and positive charged residues showing tau as having a compact conformation.

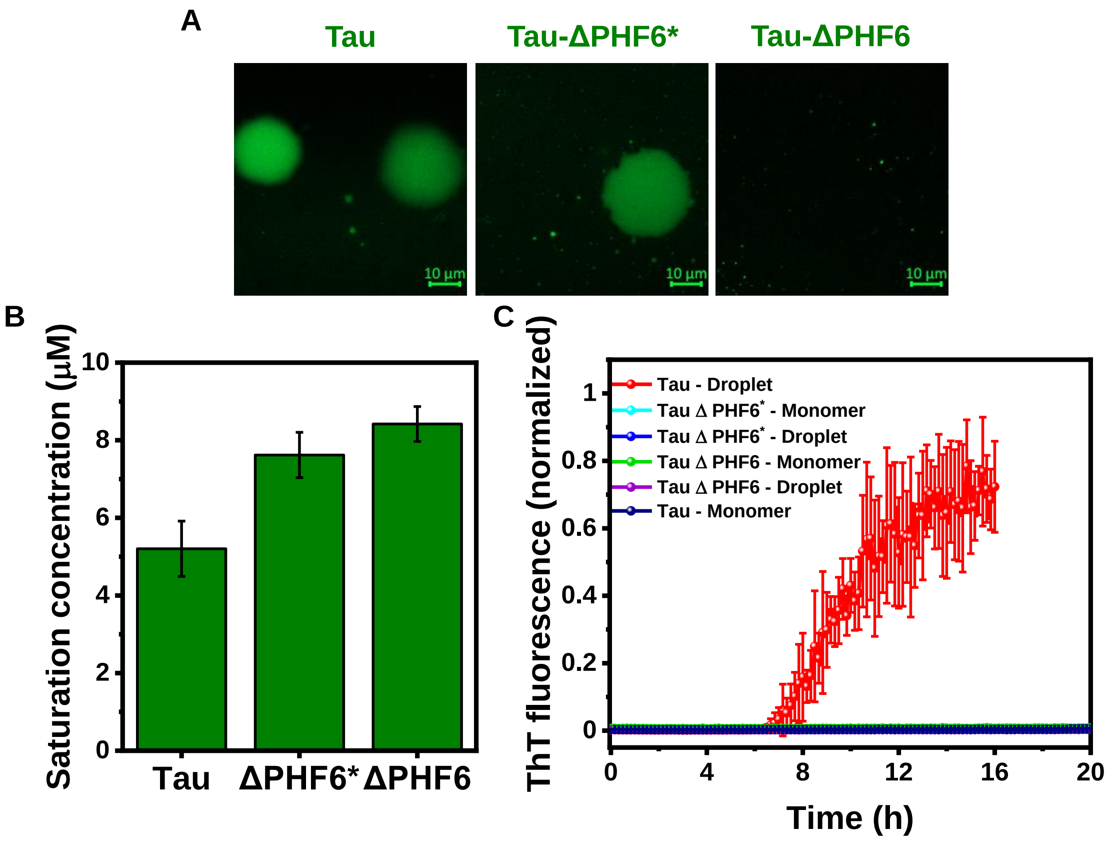


**Fig. S7. Significance of tau PHF6 in governing its phase behavior.** (*A*) Airyscan confocal images of droplets of tau, tau ΔPHF6, and tau ΔPHF6* (10 µM unlabeled protein was doped with 1% tau-Q244C-Alexa Fluor 488; green. Scale bar, 10 µm). The image for tau-only droplets is same as shown in Fig.1D for comparison. (*B*) The saturation concentration ($C_{sat}$) of tau for tau, tau ΔPHF6, and tau ΔPHF6* condensates is estimated by high-speed centrifugation. Data represent mean ± SD; n =3 independent reactions. (*C*) ThT kinetics of phase separation-mediated aggregation of tau via a liquid-to-solid transition; and separately, monomeric tau ΔPHF6 and droplet, monomeric tau ΔPHF6* and droplet, and monomeric full-length tau. The same kinetics as in (Fig. 5*A*) have been used in the case of tau 10 µM for comparison. Wherever used, full-length and tau deletion variant concentrations were kept constant at 10 µM. The data represent mean ± SD; n = 3 independent experiments.
